# SpliceSelectNet: A Hierarchical Transformer-Based Deep Learning Model for Splice Site Prediction

**DOI:** 10.1101/2025.02.17.638749

**Authors:** Yuna Miyachi, Kenta Nakai

## Abstract

Accurate RNA splicing is essential for gene expression and protein function, yet the mechanisms governing splice site recognition remain incompletely understood. Aberrant splicing caused by mutations can lead to severe diseases, including cancer and genetic disorders, underscoring the need for accurate computational tools to predict splice sites and detect disruptions. Existing methods have made significant advances in splice site prediction but are often limited in handling long-range dependencies due to high computational costs, a factor critical to splicing regulation. Moreover, many models lack interpretability, hindering efforts to elucidate the underlying biological mechanisms. Here, we present SpliceSelectNet (SSNet), a hierarchical Transformer-based deep learning model that predicts splice sites from DNA sequences spanning up to 100 kb. By integrating local and global attention mechanisms, SSNet efficiently captures both proximal and distal regulatory signals while maintaining single-nucleotide resolution. Across multiple benchmark datasets, SSNet achieves state-of-the-art performance in splice site prediction and aberrant splicing detection. Systematic in-silico mutagenesis demonstrates that attention scores reflect functional sequence importance, supporting their biological relevance. Long-range sequence perturbation experiments further show that SSNet captures distal regulatory effects beyond conventional receptive fields. Together, these results establish SSNet as a biologically interpretable framework for modeling long-range splicing regulation from genomic sequence.

## Introduction

RNA splicing is a fundamental process in eukaryotic gene expression that allows for the removal of non-coding introns and the joining of coding exons to generate mature mRNA. This intricate mechanism is essential for producing functional proteins and is tightly regulated by various cis-acting elements and transacting factors. Misregulation of splicing can lead to aberrant splicing events, which are involved in numerous diseases, including multiple types of cancer [1], neurodegenerative disorders [2], and genetic syndromes [3]. For example, in the BRCA1 and BRCA2 genes, which are known to be associated with breast and ovarian cancer, aberrant alternative splicing is highly associated with the development of breast cancer [4]. As an additional example, Duchenne muscular dystrophy is also caused by a variant in the dystrophin gene [5]. This variant leads to exon skipping, which induces a frameshift mutation and results in the loss of functions essential for maintaining muscle cell membrane integrity. In response to this, exon-skipping drugs have been developed to avoid frameshift by skipping neighboring exons, resulting in the production of a protein that is shorter than normal but still retains its function [6]. Consequently, understanding and predicting splicing is critical for both basic research and clinical applications.

To better understand splicing events, comprehensive databases and systematic analyzes have been developed. One of the first examples is the creation of a database of aberrant splicing mutations in human genes [7]. This database has provided a foundation for identifying common patterns in splicing errors and understanding their mechanisms. Building upon such foundational work, researchers have developed computational tools to analyze and predict splicing events. For example, tools such as MaxEntScan [8] utilize motif-based approaches and sequence-based statistical models to predict splice site strength. This method relies heavily on local sequence dependencies and statistical assumptions, which can limit their accuracy and flexibility when dealing with complex genomic variations.

Recent advances in computational methods for the prediction of splice sites have leveraged machine learning and deep learning techniques to enhance predictive accuracy. The advent of deep learning has provided a new avenue for addressing these challenges, allowing models to learn complex patterns from large datasets.

Among the notable approaches in this field is SpliceAI [9], a deep learning model that utilizes a neural network architecture to predict splice sites based solely on nucleotide sequences. SpliceAI has set new benchmarks in splice site prediction accuracy; however, it only consists of convolutional layers and may lack high interpretability without post-hoc analyses such as in silico mutagenesis or gradient-based attribution methods. Another prominent tool is Pangolin [10], which integrates RNA-seq data from multiple tissues to improve splice site predictions. It employs a similar deep learning framework but emphasizes splice site usage rates rather than binary classification, providing a more nuanced view of splice site strength. Although Pangolin has shown improvements over previous models, it does not explicitly address the interpretability of its predictions.

DNABERT [11], a variant of the BERT [12] model adapted for DNA sequences, has also been introduced to the field. In recent years, SpliceBERT [13] has been proposed as a BERT model pre-trained on mRNA precursor sequences from numerous biological species and fine-tuned using splice site prediction. By leveraging the transformer architecture, SpliceBERT captures contextual relationships between sequences, allowing the model to perform various genomic tasks, including splice site prediction. However, like many existing models, SpliceBERT primarily focuses on sequence representation without specialized mechanisms for handling the long-range dependencies inherent in splicing. Since SpliceBERT just adopted the same architecture as BERT, used in natural language processing, its receptive field is just 900 nt, which is too short comapred with CNN models such as SpliceAI. Regulatory elements like splicing enhancers and silencers are located far from splice sites and play a critical role in determining the splicing outcome. In the scenario of aberrant splicing, intronic mutations several kilobases away from canonical splice sites can activate cryptic splice sites or disrupt regulatory motifs, causing aberrant splicing. Thus, considering long-range interactions is essential for accurate splice site prediction. In addition, local cis-regulatory elements such as exonic splicing enhancers (ESEs) and intronic splicing enhancers (ISEs) are also key determinants of splice site usage. However, existing computational models rarely capture their contributions explicitly, limiting their ability to provide mechanistic insights into splicing regulation. Some other Transformer-based models for splice site prediction like Spliceformer [14] and SpliceTransformer [15] can incorporate longer distance relationships than SpliceBERT, but often suffer from high computational costs of the attention mechanism and lack efficiency in the inference time.

To address these issues, we propose SpliceSelectNet (SSNet), a hierarchical Transformer [16]-based model designed specifically for splice site prediction from DNA sequences. SSNet uniquely integrates local and global attention mechanisms to capture long-range dependencies (up to 100 kb) while maintaining computational efficiency. Our proposed model also directly provides attention heatmaps that highlight important regions for splice site prediction.

This model aims not only to accurately predict splice sites, but also to identify aberrant splicing events caused by mutations, classifying them as either pathogenic or benign.

To enhance scalability for long sequences, various Transformer variants have been proposed to reduce computational complexity. These include sparse attention mechanisms like Longformer [17], which limit the attention field to local windows to achieve linear scaling. However, for splice site prediction, pre-defined sparse patterns may be insufficient to capture the dense and complex interactions across distal genomic regions. Other designs, such as the Funnel-Transformer [18] and Hourglass Transformer [19], introduce hierarchical structures but typically initiate computation with full-resolution attention on the entire input sequence. For genomic tasks involving 100-kb contexts, this initial full-resolution operation remains computationally impractical.

To overcome these limitations, we adopt a local-to-global hierarchical approach inspired by the Swin Transformer [20]. By partitioning the 100-kb input into local patches and progressively merging them into higher-level representations, this strategy avoids the initial quadratic cost of long-range attention while enabling dense attention at each stage of the hierarchy. SSNet adapts this hierarchical merging process to DNA sequences by employing position-specific linear transformations on flattened local blocks. This architecture provides a practical framework for 1-bp resolution splice site prediction across 100-kb genomic contexts.

The objectives of this study are twofold: (1) to develop a highly accurate, interpretable, and efficient model for splice site prediction using dense attention mechanisms and (2) to evaluate the model’s performance in predicting aberrant splicing in disease-related datasets. By addressing the limitations of existing models and providing insight into splicing mechanisms, we aim to contribute valuable knowledge to the field of genomics and personalized medicine.

## Methods

### Training Data

For this study, we utilized three primary datasets for training: Gencode, GTEx, and the dataset used in [10] (hereafter referred to as the Pangolin dataset), each contributing unique features for splice site prediction. We downloaded the DNA sequence hg19 assembly from the UCSC Genome Browser [21], and a DNA sequence between the start and end sites of the transcript was extracted for each gene. We finally padded examples shorter than the input length by zero value and split examples longer than the input length to align with it. When dividing an example, we split it sequentially from front to back, ensuring the last part matched the input length with overlap allowed. To split this dataset into one for training and evaluation, we used the genes in chromosomes 2, 4, 6, 8, 10-22, X, and Y for training, and the genes in chromosomes 1, 3, 5, 7, and 9 that have no paralogs for evaluation. The exclusion of paralogs was performed following the criteria described by Jaganathan et al [9].

### Gencode Dataset

This dataset is based on the Gencode V24lift37 gene annotations [22]. First, only protein-coding genes were extracted from the database. There were multiple isoforms for one gene, therefore a single principal transcript was used to make the acceptor/donor label. We removed transcripts that consist of only one exon, such that any transcript in this dataset has at least one intron. Using exon region information from Gencode annotations, we defined acceptor/donor sites as the first/last nucleotide position of each exon, excluding the first/last exon, respectively. We assigned a non-splice site label for the nucleotide position that was neither acceptor nor donor site. This process assigned each nucleotide an acceptor, donor, or non-splice site label. This dataset is identical to that used in SpliceAI, but this study also used additional exon/intron labels.

### GTEx Dataset

To enhance our model’s ability to predict alternative splice sites, we incorporated additional splice sites from the GTEx dataset [23, 24], which includes intron junction information. Possible splice sites were identified from the end position of each junction seen in more than four individuals, and we ignored the end position beyond the ends of the transcript used in the Gencode dataset. The Gencode dataset primarily contains constitutive splice sites, while the GTEx dataset provides alternative splice sites. This integration resulted in a total number of splice sites that is approximately 1.5 times greater than the Gencode dataset alone. This dataset is also used in SpliceAI. As for exon/intron labels, the same labels as the Gencode dataet were used.

### Pangolin Dataset

We used RNA-seq data from seven human tissues: brain, cerebellum, heart, kidney, liver, ovary, and testis [25]. This dataset was generated from RNA-seq data used in Pangolin. First, RNA-seq data were mapped to their genomes using STAR 2.7.11 [26] based on Gencode V24lift37 gene annotations. Next, we assigned multi-mapped reads to a single location using MMR [27]. Instead of using binary labels (0/1), we calculated the splice site usage rates (continuous values between 0 and 1) as labels. Splice-Site Strength Estimate (SSE) values were calculated using SpliSER 0.1.7 [28]. While Pangolin calculated SSE for each tissue, this study did not make a distinction and used the average value of each tissue as the label. Notably, we did not utilize exon/intron labels for training on this dataset. This decision was based on two factors. First, applying exon/intron labels directly from annotations resulted in inconsistencies with the donor and acceptor site labels calculated from RNA-seq data. Second, although we attempted to incorporate Percent Spliced In (PSI) values derived from RNA-seq, our experiments showed that the model achieved superior performance without them. A summary of the training datasets is provided in Supplementary Table S1.

### Model Architecture

Our SSNet model is built on a hierarchical Transformer architecture designed to efficiently capture both local and global dependencies in nucleotide sequences. The architecture consists of the following components (Figure 1 a):

- Convolutional Layer: This layer extracts local features, allowing the model to recognize short-range interactions crucial for splice site identification, such as the GT-AG rule.
- Local Attention Mechanism: This layer is a novel aspect of our proposed architecture. It is designed to focus on nearby nucleotides (within 160bp), providing high-resolution attention to regions essential for donor and acceptor site recognition. By prioritizing the local context, the model can identify patterns and signals that indicate the presence of splice sites, enhancing prediction accuracy near the target sites.
- Global Attention Mechanism: Complementing the local attention, the global attention layer captures dependencies over long distances (up to 100 kb). This is particularly important for splicing regulation, as regulatory elements can be located far from the splice sites they influence. The integration of global attention allows the model to incorporate distant interactions and contextual information, which traditional models often overlook.

**Fig. 1:**
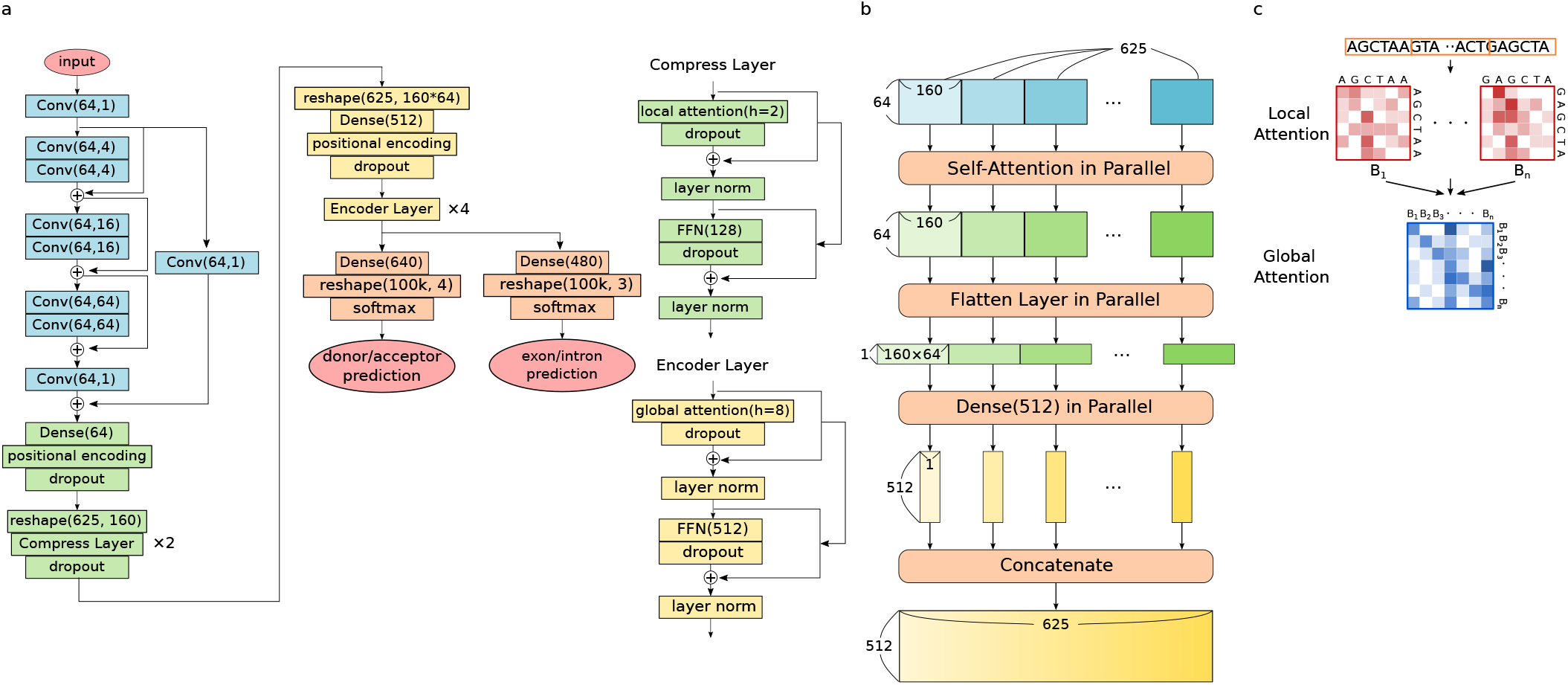
(a) The overall architecture of SSNet. It consists of convolutional layers (blue), local attention layers (green), and global attention layers (yellow). (b) The detailed architecture of local attention layers. (c) The concept of hierarchical attention mechanism.

First, input sequences are transformed to a matrix by one-hot encoding, A, C, G, T, and padding mapped to [1, 0, 0, 0], [0, 1, 0, 0], [0, 0, 1, 0], [0, 0, 0, 1], and [0, 0, 0, 0]. Second, the encoded matrix is fed to the convolutional block with a residual connection. In Figure 1 a, Conv(N, W) represents a convolutional layer, where N is the number of kernels and W is the kernel size. Third, the extracted local information through convolutional blocks is handled in the local attention block. The input matrix is compressed so that the global attention mechanism can handle it at once. The process of computation in the local attention mechanism is explained in detail in Section Local Attention. Then, the compressed information is fed to the global attention block, and the model learns an end-to-end global mutual relationship. The global attention mechanism is the multihead self-attention mechanism with 8 heads. Finally, the global information is fed to each dense layer, reshaped to length 100k, and activated with a softmax function to predict the possibility of an acceptor/donor/non-splice site and that of an exon/intron for each nucleotide position. The loss value is calculated for each prediction and the sum of them is used for optimization. The detailed loss function is explained in Section Loss Function.

In the conventional attention mechanism, the computational complexity is the square of the input length, so the limit of dense attention is a few hundred tokens. In this architecture, a two-stage hierarchical attention mechanism is combined to reduce the input length to a few hundred tokens in a single attention computation, thus preserving computational efficiency while learning long-range array interactions in the model as a whole. Figure 1 c illustrates the concept of the hierarchical attention mechanism. In local attention, the model divides the input into blocks consisting of short sequences and calculates the interactions within those blocks. Then, in global attention, the model computes the interaction between each block.

While sparse attention, such as linear attention [29], is often used to reduce the computational complexity to a polynomial input length to handle long-range sequences in the attention mechanism, this study proposes a new method to handle long-range sequences while maintaining dense attention. The dense matrix of local attention and global attention weights makes it possible to visualize which regions the model focuses on for prediction, thus preserving high interpretability.

### Local Attention

Figure 1 b illustrates a process of local attention in SSNet-100k. First, input matrices that have a length of 100k are divided into 625 blocks with a length of 160 nt. Next, each block is fed to multi-head self-attention mechanisms with 2 heads using relative positional encoding [30, 31] in parallel. The output of each mechanism is of the same shape as the input, 160 nt × 64. Through a flatten layer and a dense layer, we have a 512-dimension vector from each output. We now have 625 vectors with 512 dimensions as a whole. Finally, we concatenate all of them and obtain an output matrix of size 625 × 512. Since the length of the matrix becomes 625 from 100k, the model can compute an end-to-end relationship in the global attention mechanism.

For models with input lengths of 1k and 10k, the first division into blocks was changed to 50 blocks of length 20 and 50 blocks of length 200, respectively. Other than that, the same parameters were used for the 100k input length model.

Let *M* = (*M*_1_, *M*_2_, …, *M*_*B*_) be the input of the local attention, where *B* is the number of blocks. Using *L* as the length, *D* as the dimension of the input, the shape of *M* is represented as *L* × *D* and *M*_*i*_ as the *B* × *L*_*B*_ × *D*. (*L*_*B*_ = *L/B*) Using *H* as the number of heads, the local attention is defined by the following equations.

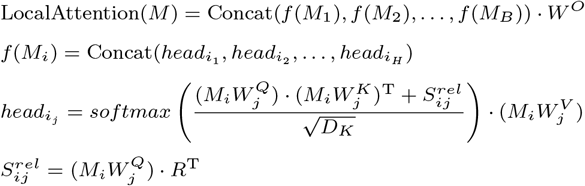

This is almost the same as the equations in Multi-Head Self-Attention. 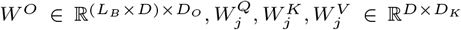 are the matrices for the linear projection. *D*_*O*_ is the output dimension and *D*_*K*_ is the output dimension of each head (*D*_*K*_ = *D/H*). 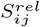 represents the relative positional embeddings interacted with queries. 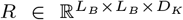 is a matrix of relative positional encodings. When use *R*_*kl*_ as the element of matrix *R, R*_*kl*_ is defined as *E*_*l*−*k*_, where 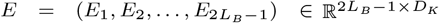. Here, *E*_*l*−*k*_ represents the relative positional encoding according to the distance from query k to query l. For the effective computation of relative positional encoding, we used the skew algorithm proposed in [31] in code implementation.

### Training Procedure

We trained four variations of the SSNet model depending on the training dataset. All models were initially trained on the Gencode dataset for 400 epochs. Variations were as follows:

- SSNet_pangolin: Trained on the Pangolin dataset for an additional 50 epochs.
- SSNet_gtex: Trained on the GTEx dataset for 50 epochs.
- SSNet_gtex_pangolin: Trained on the GTEx dataset followed by the Pangolin dataset, each for 50 epochs.
- SSNet_pangolin_gtex: Trained on the Pangolin dataset followed by the GTEx dataset, each for 50 epochs.

Because the performance trend was largely determined by the dataset used in the final training step, SSNet_gtex and SSNet_pangolin_gtex showed similar performance (both ending with GTEx), and likewise SSNet_pangolin and SSNet_gtex_pangolin showed comparable results (both ending with Pangolin). For clarity, we therefore focus on the two-step training models (SSNet_pangolin_gtex and SSNet_gtex_pangolin) in the performance comparison. These two variants not only represent the performance tendencies of the single-dataset models but also demonstrated more stable results across benchmarks.

We trained the model 5 times for each variation, randomly dividing the training datasets into 90% training dataset and 10% validation dataset. The validation data was used for hyperparameter optimization and early stopping, which was to stop the training when it converges. We used the Tensorflow library and Adam as the optimization method. The training was performed using Tesla V100, NVIDIA A100, and NVIDIA H100 with a batch size of 16. The learning rate was set to warm up linearly to 0.00005 in the first 80000 steps and then decreased linearly to 0.0 by 800000 steps. Mixed precision, using both 16-bit and 32-bit floating-point types, was used to train the model faster and use less memory. However, for the part that uses softmax as an activation function, we used a 32-bit floating-point type because softmax is a numerically unstable computation.

The source code and the trained models are publicly available at https://github.com/yuna728/SSNet.

### Loss Function

Cross entropy was used as the loss function. Let *y* be the correct answer labels represented by categorical variables and *p* the probability distribution of output. Suppose that the correct label *y* takes the value *i*. We denote the corresponding *i*-th probability *p*_*i*_. We define *p*_*t*_ as the probability assigned to the correct label. Cross entropy loss *CE*(*p*_*t*_) is defined as follows;

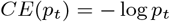

Our dataset was imbalanced because the number of acceptor/donor sites was much smaller than that of non-splice sites (donor/acceptor site labels only accounted for 0.033% on the Gencode dataset) and the length of exon was much shorter than that of intron (exon labels accounted for 5.82% on the Gencode dataset). Thus, to improve that imbalanced situation, we combined balanced cross entropy and focal loss [32].

Balanced cross entropy *BCE*(*p*_*t*_) was defined in the following equation. By introducing *α* and setting larger values to fewer labels, the impact of fewer label examples on loss values increases. Suppose that the value of the correct answer label *y* was *i*, let the *i*-th *α* value *α*_*i*_ be denoted by *α*_*t*_.

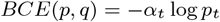

Focal loss function *FL*(*p*_*t*_) was represented as follows. By introducing *γ* ≥ 0, the loss value for more difficult examples accounts for a larger proportion of the total loss.

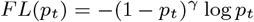

When *p*_*t*_ is close to 0, which is an example where the difference between the output and the correct label is large, and the model is not well-trained, 1 − *p*_*t*_ is close to 1, so the effect of introducing *γ* on the loss is small. On the other hand, when *p*_*t*_ is close to 1, this is when the model can produce an output close to the correct label, and in this case, 1 − *p*_*t*_ is small. Therefore, where *γ >* 0, the loss becomes smaller compared to the standard cross-entropy without introducing *γ*.

Combining these two loss functions defined by *α* and *γ*, the loss function used in training *BFL*(*p*_*t*_) was defined as follows:

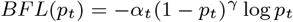

For the *α* value, we used 8.0/8.0/3.0/1.0 for donor/acceptor/non-splice/padding labels and 0.7/0.25/0.05 for exon/intron/padding labels in the training on the Gencode Dataset. For additional learning on the GTEx dataset, 6.0/6.0/5.0/3.0 for donor/acceptor/non-splice/padding labels and 0.7/0.25/0.05 for exon/intron/padding labels were used. For additional learning on the Pangolin dataset, 8.0/8.0/3.0/1.0 for donor/acceptor/non-splice/padding and 0.0 for all exon/intron/padding labels were used. As for the *γ* value, we used 2.0 in both phases of training because it showed the best result in [32].

### Validation Dataset

For model validation, we mainly utilized three datasets:

- SpliceVarDB Dataset: This is an online database of over 50,000 variants collected from over 500 publicly available data sources with experimentally confirmed effects on splicing. Each variant is classified as “splice-altering” (∼25%), “not splice-altering” (∼25%), or “low frequency splice-altering (∼50%) [33]. Detailed statistics of the SpliceVarDB dataset are summarized in Supplementary Table S2.
- SSCVDB Dataset: This database is a catalog of 30,130 splice-site generating mutations identified from 322,072 transcriptomes in the Sequence Read Archive by a novel information analysis method (juncmut) [34]. It does not contain negative samples (not splice-altering variants).
- BRCA Dataset: This dataset includes variants in the BRCA1 and BRCA2 genes, which are known to be involved in breast cancer. Each variant has pathogenicity label. Data were sourced from BRCA Exchange [35]. A summary of the BRCA dataset is shown in Supplementary Table S3.

For all three validation datasets, we used GRCh38/hg38 reference genome. To evaluate the performance of models, the difference between the model’s predictions for reference sequences without a mutation and those for alt sequences with a mutation was taken, and if the maximum difference was greater than a threshold value, the variant was judged splice-altering/pathogenic, otherwise not splice-altering/benign. A threshold value of 0.2 was used for all models. Confidence intervals were estimated using nonparametric bootstrap resampling. Unless otherwise noted, resampling was performed at the variant level with replacement for 1,000 iterations. For classification metrics, stratified bootstrap resampling was applied to preserve the original class distribution. For ROC and precision–recall curves, mean curves and pointwise 95% confidence intervals were obtained by interpolating bootstrap curves on a common grid. For performance comparison, we made predictions of SpliceAI [9], Pangolin [10], Spliceformer [14], SpliceTransformer [15], and SpliceBERT [13].

## Results

### Performance on Gencode and long non-coding RNAs

First, we evaluated the performance of SSNet on the Gencode protein-coding dataset and compared it with SpliceAI. Gencode protein-coding genes represent the most standard and reliable evaluation setting for splice site prediction, and also form a crucial foundation for downstream mutation effect prediction and functional analysis. Furthermore, since both SpliceAI and SSNet are models trained using Gencode annotations, evaluation using this test dataset constitutes a fundamental assessment. SSNet_base is a variant of SSNet trained solely on the Gencode dataset.

As summarized in Table 1, SSNet consistently outperforms SpliceAI in accuracy, F1 score, and Top-k accuracy, demonstrating more precise splice site localization while suppressing false detections. The improvement in precision, in particular, signifies a reduction in false positives, which is advantageous in practical scenarios where splice sites must be identified from multiple candidates. Furthermore, recall is maintained at a level comparable to SpliceAI, indicating that the performance improvement of SSNet is not achieved at the expense of sensitivity but rather through an enhanced balance between sensitivity and specificity.

**Table 1.** Performance comparison on the Gencode test dataset. Values are reported as mean (95% confidence interval), estimated using gene-level bootstrap resampling.

| Model | Recall | Precision | F1 | Top-k | auPRC |
| --- | --- | --- | --- | --- | --- |
| SpliceAI | **0.944 (0.938-0.949)** | 0.857 (0.850-0.863) | 0.898 (0.893-0.903) | 0.928 (0.923-0.932) | **0.945 (0.938-0.952)** |
| SSNet_base | 0.937 (0.932-0.941) | **0.934 (0.929-0.938)** | **0.935 (0.931-0.939)** | **0.940 (0.936-0.944)** | 0.943 (0.935-0.950) |

Furthermore, we evaluated the model’s generalization ability using a long non-coding RNA (lncRNA) dataset (Table 2) not included in the training data. We used the lncRNA dataset provided in [9]. In this dataset, SSNet achieved higher recall than SpliceAI, while SpliceAI achieved higher precision, F1 score, Top-k accuracy, and auPRC.

**Table 2.** Performance comparison on the long non-coding RNA dataset. Values are reported as mean (95% confidence interval), estimated using gene-level bootstrap resampling.

| Model | Recall | Precision | F1 | Top-k | auPRC |
| --- | --- | --- | --- | --- | --- |
| SpliceAI | 0.795 (0.771-0.816) | **0.912 (0.897-0.926)** | **0.848 (0.834-0.863)** | **0.920 (0.907-0.931)** | **0.963 (0.956-0.970)** |
| SSNet_base | **0.824 (0.805-0.845)** | 0.842 (0.825-0.859) | 0.833 (0.818-0.847) | 0.867 (0.850-0.883) | 0.927 (0.916-0.937) |

SpliceAI, trained on protein-coding genes, tends to prioritize auxiliary signals like Exonic Splicing Enhancers (ESEs) that enhance splicing efficiency. However, lncRNAs lack ESEs that bind to SR proteins promoting splicing reactions [36], leading SpliceAI to fail to capture signals at many splice sites and resulting in a significant drop in recall. Conversely, the higher Precision likely resulted from detecting only highly reliable sites possessing consensus sequences strong enough.

In contrast, SSNet maintained high Recall because, as later demonstrated by motif analysis in this study (in Quantitative Analysis of Attention by In-Silico Mutagenesis), this model specifically directs attention toward U-rich polypyridine tracts (Py-tracts). Since it is known that splicing in lncRNAs is promoted by the number of uridines in Py-tracts [36], SSNet was able to identify splice sites based on this strong signal.

However, we observed a noticeable drop in Precision for SSNet on this dataset. This performance discrepancy is further illustrated by the Precision-Recall curves provided in Supplementary Figure S1. The primary cause of this increased false-positive rate likely stems from a confounding effect introduced during model training. Because SSNet explicitly incorporates exon/intron structural annotations into its learning scheme, the model inherently captures sequence features unique to coding regions, such as codon bias, as auxiliary predictive cues. Since lncRNAs lack these coding-specific signatures, the model falls back on potential, yet non-functional, sequence motifs. Given that splicing efficiency in lncRNAs is generally low and many unfunctional decoy sites exist [37], SSNet sensitively detected even these potential sites, which reduced its precision.

Overall, the performance decline in lncRNAs is largely attributable to these non-coding-specific splicing characteristics and the model’s sensitivity to structural motifs that do not necessarily function in vivo. While this indicates a clear limitation of SSNet in distinguishing functional splice sites from non-coding decoy sites, the primary target of this model remains protein-coding genes. Given that SSNet has already demonstrated high precision and robust recall for coding regions, these characteristics do not compromise the model’s practical utility in clinical and medical genetics focused on protein-coding variants.

### Competitive Performance on SpliceVarDB

First, we evaluated aberrant splicing prediction performance on SpliceVarDB. This benchmark is a large dataset, comprising variants from over 8,000 human genes, and it specifically labels whether a variant alters splicing rather than indicating pathogenicity.

Figure 2 presents the AUROC values for all Variants (All) as well as for each variant location category (Exon, Splice Site, Intron). Across all categories, our proposed model demonstrates performance comparable to state-of-the-art methods. Notably, it consistently outperforms Spliceformer and SpliceTransformer, which are also based on Transformer architectures. Variability in AUROC is observed only in the Splice Site category, likely due to the unbalanced nature of the dataset, as most variants in this category are splice-altering. Nevertheless, all models achieve AUPRC values of 0.99 or higher, indicating highly accurate predictions in this category. Detailed comparison including additional SSNet variants is shown in Supplementary Figure S2. While the overall differences among models were relatively small, SSNet variants consistently achieved stable performance across regions. Numerical results for SpliceVarDB evaluation on each region are provided in Supplementary Table S4, confirming that SSNet models perform comparably to SpliceAI and Pangolin across All, Exon, Splice Site, and Intron categories.

**Fig. 2:**
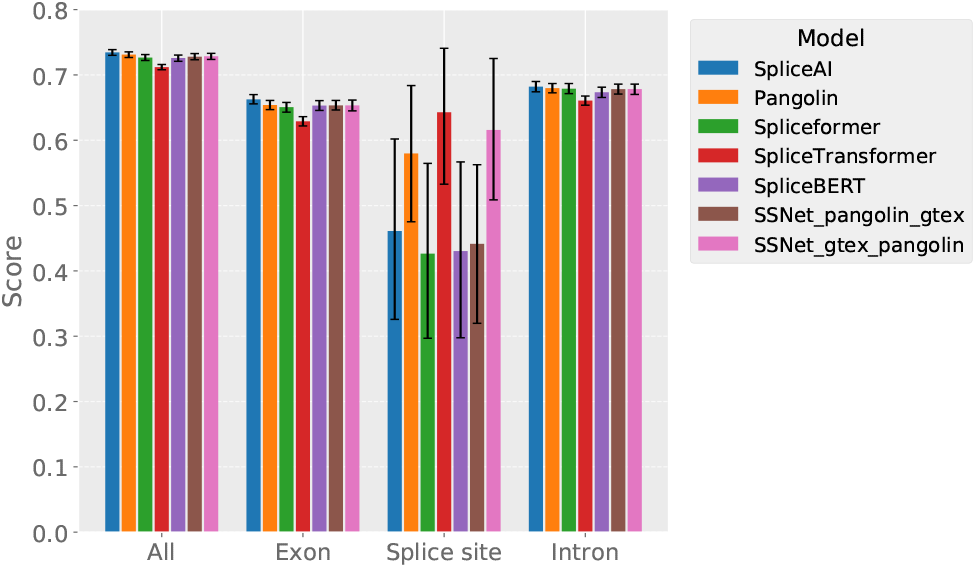
AUROC values for all variants in SpliceVarDB (All) and stratified by variant location (Exon, Splice Site, Intron). SSNet is compared with state-of-the-art methods, SpliceAI, Pangolin, Spliceformer, and SpliceTransformer. Error bars indicate 95% confidence intervals estimated by stratified bootstrap resampling.

These results indicate that SSNet achieves performance comparable to the current state-of-the-art in aberrant splicing prediction. Its robustness is further supported by its consistent performance across a wide range of genes, rather than being limited to specific loci. Furthermore, SSNet exhibits improved performance relative to other Transformer-based models.

### Stronger Sensitivity on SSCVDB

Since SSCVDB does not contain negative samples in which splicing changes do not occur, we evaluated the proportion of splicing alterations detected by each model across different thresholds. The dataset provides information on splice site positions in the original sequence (Hijacked_SS) as well as newly generated splice sites caused by mutations (Primary_SS). Accordingly, in addition to assessing the maximum change in predicted values across the entire sequence, we specifically examined changes at Hijacked_SS and Primary_SS. Thresholds were varied from 0.0 to 0.2, with 0.2 chosen as the upper limit because it is commonly used as a threshold in models such as SpliceAI. For each threshold, the proportion of predictions exceeding the threshold was plotted (Figure 3). For both the entire sequence and Primary_SS, the SSNet models trained on the GTEx dataset at last (SSNet_pangolin_gtex) achieved the highest AUC values (0.818–0.819 for the entire sequence and 0.784–0.785 for Primary_SS). This indicates that the models trained on GTEx exhibit strong sensitivity to the generation of novel splice sites. Although SpliceAI was trained on the same dataset, it achieved lower AUC values (0.722 for the entire sequence and 0.670 for Primart_SS), whereas the SSNet models consistently demonstrated significantly higher sensitivity, highlighting the superior performance of our proposed approach. Regarding Hijacked_SS, since SSCVDB is a collection of data that creates new splice sites rather than disrupts existing ones, all models showed lower detection rates compared to Primary_SS.

**Fig. 3:**
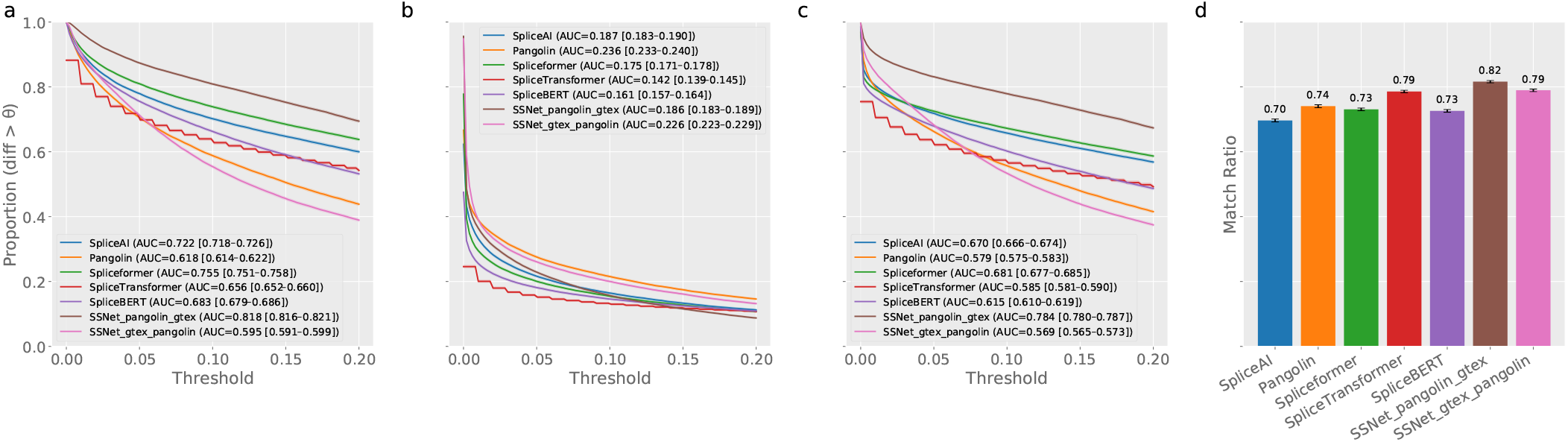
Proportion of predicted splicing changes exceeding thresholds (0.0–0.2) is shown for (a) the entire sequence, (b) original splice sites (Hijacked_SS), and (c) newly created splice sites (Primary_SS) in SSCVDB. (d) The probability that the maximum predicted change occurs at Primary_SS. Error bars indicate 95% confidence intervals estimated by stratified bootstrap resampling.

We further calculated the proportion of cases where the change in predicted values at Primary_SS corresponded to the maximum change across the entire sequence. Once again, SSNet_pangolin_gtex showed the highest match rates. This observation reflects the underlying nature of SSCVDB variants, which generate new splice sites, and further confirms the strong predictive performance of the proposed model for this dataset. Supplementary Figure S3 provides the detailed detection results for all SSNet variants on SSCVDB. SSNet_gtex showed a performance trend similar to SSNet_pangolin_gtex, both representing the best aberrant splicing detection performance among the compared models.

We acknowledge that the SSCVDB dataset primarily comprises variants that create novel splice sites, representing a relatively rare form of aberrant splicing. To further assess the model’s robustness across different splicing patterns, we also evaluated performance on the IRAVDB dataset [38]. This dataset contains only variants that cause intron retention. We calculated the predicted changes due to the variant for the splice site directly disrupted by the variant (target_pos), its partner splice site (partner_pos), and the entire input sequence (Whole_Seq) (Supplementary Figure S4).

For all models, changes in Whole_Seq and target_pos were captured with relatively high accuracy. Changes in the distant partner_pos were an indirect effect, making it an extremely difficult task that was barely captured. SSNet_pangolin_gtex demonstrated high performance with an AUC of 0.951 for Whole_Seq, comparable to SpliceAI. SSNet_gtex_pangolin achieved an AUC of 0.883 for target_pos, second only to Pangolin and nearly equivalent to SpliceAI. Furthermore, for partner_pos, although the overall values are very low, SSNet_gtex_pangolin was able to capture changes, following SpliceAI.

### Superior Performance on BRCA Dataset and Attention Analysis

We evaluated the performance of aberrant splicing prediction on the BRCA dataset. First of all, we made predictions only for variants labeled by experts (4909 positive, 2504 negative). Since the F1 score and accuracy are greatly affected by the threshold settings, we mainly look at AUROC and AUPRC to compare the performance. Figure 4 shows that the proposed model outperforms SpliceAI and Pangolin by a wide margin. In particular, SSNet_gtex_pangolin consistently showed the best performance. Second, we included predictions that were “Not Yet Reviewed” in the expert labeling but were labeled in other databases (7959 positive, 6566 negative). In this case, as before, SSNet outperforms the two state-of-the-art models in AUROC and AUPRC. In particular, SSNet_gtex_pangolin continued to perform the best. Comparing SpliceAI and Pangolin, which have the same architecture, Pangolin shows higher performance in both cases, and SSNet_gtex_pangolin shows the best performance when compared within SSNet variations. The model trained on the pangolin dataset, which takes into account the strength of the splice sites, tends to perform better than the other models that were not trained on the pangolin dataset. Supplementary Figure S5 provides ROC and PR curves for all models on the BRCA dataset. SSNet_pangolin showed performance comparable to SSNet_gtex_pangolin, both achieving high AUROC and AUPRC values due to incorporating splice-site strength in training. Supplementary Table S5 reports the detailed evaluation results on the BRCA dataset, further supporting the advantage of Pangolin-trained SSNet variants in this benchmark.

**Fig. 4:**
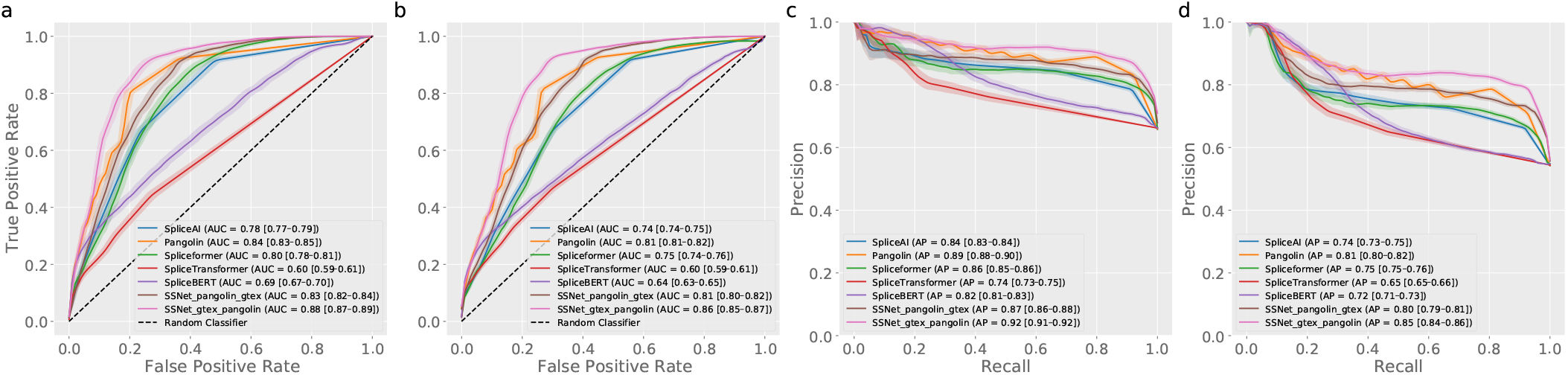
Performance of SpliceAI, Pangolin, Spliceformer, SpliceTransformer and SSNet for BRCA dataset. Shaded regions indicate 95% confidence intervals estimated by stratified bootstrap resampling. (a) ROC curves for expert label. (b) ROC curves for all label. (c) PR curves for expert label. (d) PR curves for all label.

In particular, SSNet_pangolin showed the best performance among the Pangolin-ending models, which was comparable to SSNet_gtex_pangolin. We therefore use SSNet_pangolin as a representative for detailed attention analysis. We also compared the distribution of predictions between SSNet_pangolin and SpliceAI, plotting only those variants present in the BRCA1 gene that were labeled pathogenic in Figure 5. The x-axis represents position, the y-axis represents predictions in SSNet_pangolin, and the color represents benign/pathogenic predictions made by SpliceAI. The blue plots show variants predicted as benign and the orange plots show one as pathogenic. Focusing on Exon 10, the largest exon in the BRCA1 gene, we see that it is correctly predicted as pathogenic by SSNet_pangolin with a predicted value of 0.2 or higher, while in SpliceAI variants predicted as benign (predicted values are 0.2 or lower) are concentrated in this exon (in the red circle of Figure 5). A similar trend was observed when comparing the predictions of SSNet_pangolin and Pangolin. It is known that Exon 10 is a region where many variants related to breast cancer pathogenicity are clustered, and it is also known to be a place where complex splicing regulation takes place in relatively long exons. Therefore, we examined in detail several pathogenic variants in Exon 10, which is above 0.2 in SSNet_pangolin and below 0.2 in SpliceAI, as described earlier.

**Fig. 5:**
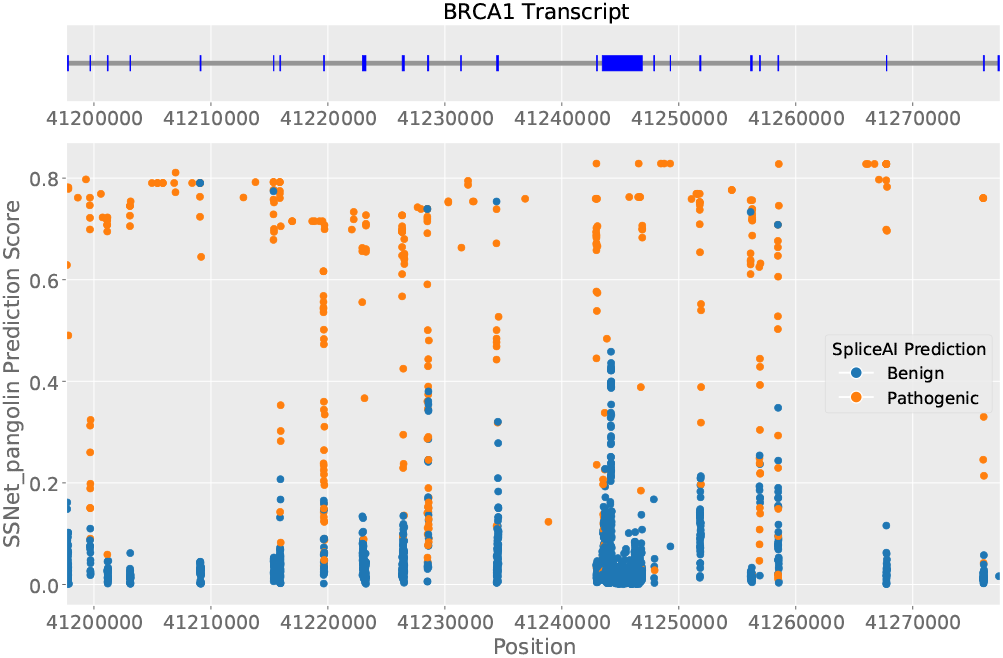
Prediction value of SSNet and SpliceAI of pathogenic variants located in BRCA1 gene. The x-axis represents position, the y-axis represents predictions in SSNet_pangolin, and the color blue/orange represents benign/pathogenic predictions made by SpliceAI. The area circled in red is the cluster of mutations in Exon 10 that have a prediction greater than 0.2 in SSNet but less than 0.2 in SpliceAI.

We found that a common cryptic acceptor site was activated in many of the variants. Figure 6 a shows the change in SSNet predictions in the chr17:g.41244217:G*>*A variant. The cryptic acceptor site in Exon 10 is a very weak splice site in Ref Seq but changes to a relatively strong splice site in Alt Seq. To examine the change in attention of this variant, we visualized the Local Attention at Block 209, where the variant and cryptic acceptor site are located, and at Block 208 before it (Figure 6 c,d). The global attention between the surrounding Blocks is also shown (Figure 6 b). Unless otherwise stated, we analyze attention maps averaged across heads and layers. We also found that specific sites upstream of the variants were activated in the attention map, even though the position of the variants varied. In this variant, we see that the location about 160 bases upstream from the variant is strongly noted in Block 208. The downstream region of the variant also shows a large change in attention, while the cryptic acceptor site itself shows a relatively small change in Block 209. In addition, focusing on global attention, the effect is noticeably stronger from Block 208, where a strongly activated region was present, to Block 209, where a cryptic acceptor site is present. This suggests that the activation region of Block 208 is involved in some kind of splicing regulation, which activates the cryptic acceptor site in the downstream region as a splice site. Although no splicing regulators have been reported for these regions in previous studies to the best of our knowledge, it is possible that the exonic splicing enhancer is activated by the variant or that the exonic splicing silencer is destroyed and the cryptic acceptor site downstream of the variant is activated.

**Fig. 6:**
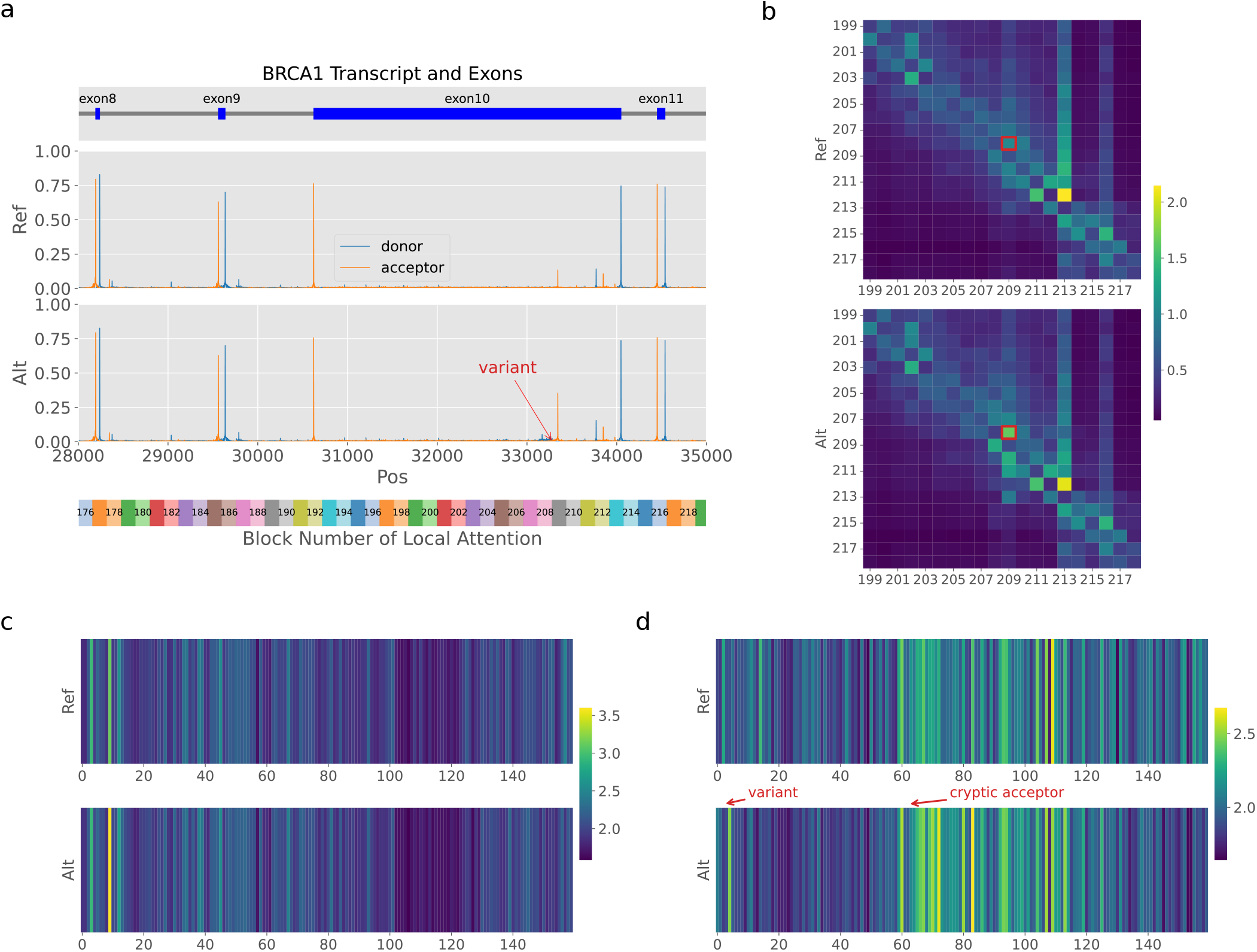
Variant chr17:g.41244217:G*>*A in the BRCA1 gene. (a) Gene structure and SSNet predictions. Top: BRCA1 gene structure; second: predicted splice site probabilities for the reference sequence; third: predictions for the alternative sequence; fourth: corresponding local attention block numbers. Donor site probabilities are shown in blue, acceptor site probabilities in orange. The variant position is indicated by a red arrow. A cryptic acceptor site (in Block 209) downstream of the variant is activated in the alternative sequence. X-axis represents the relative distance from the start of the BRCA1 transcript. (b–d) Attention weight heatmaps for the variant. Top: reference sequence; bottom: alternative sequence. (b) Global attention around the variant, indicating increased interaction from Block 208 to Block 209 upon mutation (red box). (c) Local attention of Block 208 (nucleotides 2498–2657 relative to Exon 10), showing a strongly activated region upstream. (d) Local attention of Block 209 (nucleotides 2658–2817 relative to Exon 10). The variant is at position 2661. A substantial change in attention weights is observed downstream of the variant.

### Attention Analysis Case Study of ESE/ISE

In the previous section, we reported the results of attention analysis in the BRCA1 gene. We do not claim attention scores to be direct mechanistic explanations. Instead, we view dense attention as an interpretable intermediate signal that highlights sequence regions influential to the model’s predictions. To further demonstrate the utility of the proposed model’s attention weights in analyzing cis-regulatory elements involved in splicing, such as exonic splicing enhancers (ESEs), we conducted a case study on the mouse IgM gene. In the M2 exon of the IgM gene, an ESE is present at the 5’ end, and it is known that this is important for the activation of the acceptor site immediately upstream [39].

To test this, we applied the SSNet_gtex_pangolin model to the IgM gene and assessed whether it could accurately predict the influence of the ESE. We compared the predictions using the wildtype IgM gene sequence with predictions where the first 42 bp at the 5^′^ end of Exon 6, corresponding to the known ESE, were replaced with “N.” In our model, because “N” is encoded as the uniform average of the four nucleotides “A,” “C,” “G,” and ‘T’ ([0.25, 0.25, 0.25, 0.25]), replacing nucleotides with “N” effectively masks the sequence. As shown in Figure 7 a, this in silico experiment demonstrates that masking the ESE abolishes the activity of the acceptor site at the 5’ end of Exon 6. We validated this result also in SpliceAI prediction (Supplementary Figure S6).

**Fig. 7:**
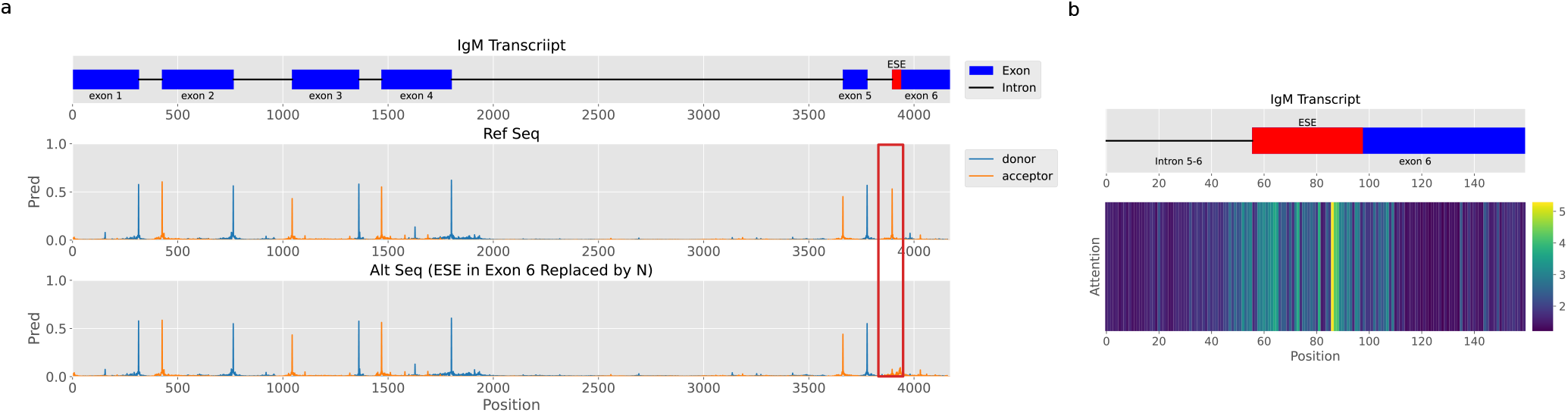
(a) SSNet_gtex_pangolin predictions for the mouse IgM gene. Top: gene structure with introns (black lines) and exons (blue boxes); the 42 bp ESE at the 5’ end of Exon 6 is highlighted in red. Bottom: predicted donor (blue) and acceptor (orange) probabilities for the wild-type sequence and with the ESE region masked with “N”. Masking the ESE abolishes the acceptor site at the 5’ end of Exon 6 (red box). (b) Local attention weights near Exon 6, showing high attention in the ESE region relative to other exon positions.

These results indicate that the proposed model not only captures the characteristics of consensus sequences in introns, such as the GT-AG rule, but also recognizes the functional characteristics of exonic splicing enhancers, accurately predicting their effects. The IgM exon 6 ESE analyzed here is a classical and well-established example, which serves as a proof-of-concept that SSNet can recover known enhancer functions. This suggests that the approach may be extended to uncover less characterized regulatory motifs in future studies.

We further visualized local attention values around Exon 6 (Figure 7b), which revealed that attention is high within the ESE region relative to other regions. This suggests that even without prior knowledge of the ESE location, the region can be inferred through analysis of attention scores.

In addition, we conducted an in silico experiment using Exon 6 of the FAS gene as an example to detect intronic splicing enhancers (ISE). An intronic splicing enhancer (ISE), URI6, is located downstream of Exon 6 in the FAS gene, and its disruption reduces Exon 6 inclusion [40]. Furthermore, it is also known that even when the ISE is disrupted, introducing a mutation downstream of Exon 6 that enhances the consensus sequence of a donor site called U1C can restore Exon 6 inclusion. Figure 8 (a) shows the positions of URI6 and U1C within the FAS gene, while Figure 8 (b) illustrates the prediction results for WT, mURI6, and mURI6+U1C using SSNet. As confirmed in in vitro experiments, SSNet predictions recapitulate these experimental observations, showing that URI6 mutations reduce the probability of upstream and downstream splice sites of Exon 6, whereas U1C mutations restore splice site strength. We validated this result also in SpliceAI prediction (Supplementary Figure S7). This indicates that SSNet can accurately predict exon skipping caused by ISE disruption and has learned the cooperative action between splicing enhancers and the strength of consensus sequences around splice sites.

**Fig. 8:**
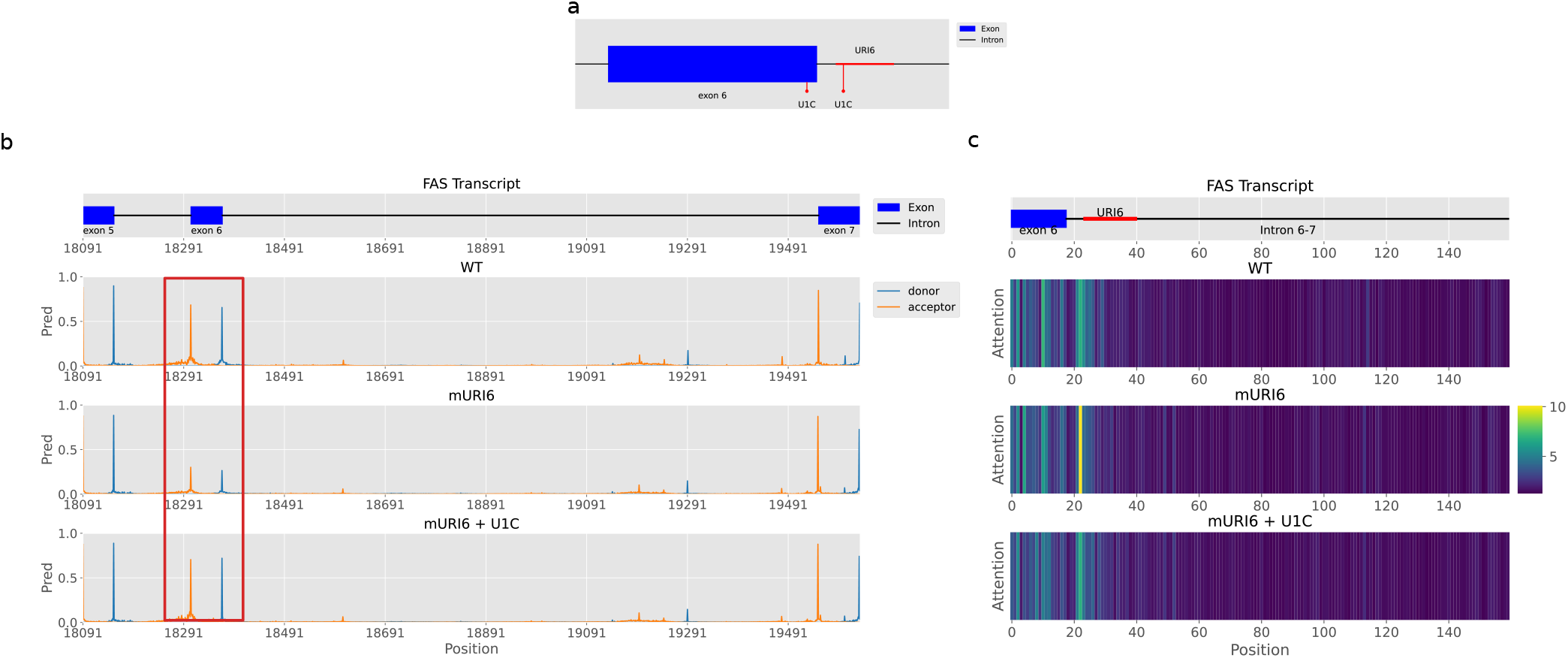
(a) Positions of intronic splicing enhancer URI6 (+7 to +23 of Intron 6) and U1C mutant sites relative to Exon 6 in the FAS gene. U1C introduces substitutions at positions -3 and -2 of the 3’ end of Exon 6 (to C and A, respectively) and positions +7 and +8 of the 5’ end of Intron 6 (to A and T, respectively). (b) SSNet predictions for wild-type, mURI6, and mURI6+U1C sequences. The red boxes indicate regions where splice site probabilities are altered by mutations. (c) Local attention weights (Block 126) near Exon 6, showing differential attention across wild-type and mutant sequences, highlighting cooperative cis-element interactions.

Furthermore, to investigate which regions within each sequence received attention, we visualized the local attention near Exon 6 (Figure 8 (c)). We note that attention scores represent a probability distribution summing to one; therefore, an increase in one region often reflects a relative redistribution of focus due to the loss of signal elsewhere. In the wild-type construct, the model assigned attention to both the downstream region of exon 6 and the 5’ region of URI6. Intriguingly, in the mURI6 mutant, while attention at the mutated URI6 site decreased, we observed a compensatory increase in attention at the region upstream of URI6. This suggests that with URI6 disrupted, the model shifts its focus intensely to remaining secondary cis-elements to maintain prediction confidence. In the U1C+mURI6 background, this hyper-focus on the upstream region diminished, and attention patterns became more balanced. This indicates that strengthening the 5^′^ splice site recognition through U1C compensates for the enhancer loss, thereby relieving the model’s reliance on the upstream auxiliary elements. Taken together, these patterns suggest that the model has learned to integrate multiple partially redundant cis-elements: an intronic splicing enhancer of URI6, a downstream auxiliary element in Exon 6, and flanking nucleotides upstream of URI6. Disruption of URI6 reduces the availability of these cooperative signals, forcing the model to rely on weaker cues, whereas strengthening the 5^′^ splice site through U1C compensates in part by restoring the contribution of other elements. Changes in attention are not restricted to the mutated position, but extend to surroundings, reflecting a context-dependent reweighting of available signals. Thus, the attention dynamics recapitulate the interplay between enhancer activity and splice-site strength, supporting the notion that the model captures context-dependent redundancy among cis-regulatory features.

### Quantitative Analysis of Attention by In-Silico Mutagenesis

In the previous section, we focused on ESE/ISE in the IgM and FAS genes as a case study for attention analysis. Although attention does not directly provide explanations for predictions, the previous section revealed correlations between high-attention regions and splicing enhancer regions. Here, we performed an in silico mutagenesis experiment in which the high-attention and low-attention regions were masked individually, and the resulting predictions were compared with those obtained using the unmasked sequence as input to quantitatively compare the impact of each region on the predictions.

Using the test dataset from the Gencode dataset, prediction and attention calculations of reference sequences were first performed. Next, for high attention regions, areas exceeding the average and at least five times the minimum value for both global and local attention were extracted. For low attention regions, areas below the average and at most three times the minimum value for both global and local attention were extracted. It is obvious that predictions change significantly when masking around splice sites, so we excluded sequences overlapping the 5-bp regions before and after the splice site. Additionally, regions at the end of sequences where padding follows tend to show high attention regardless of function, so we excluded these as well. To ensure the same number of motifs were masked for each input sequence, we randomly selected masking positions within candidate regions for low-attention regions to match the number in high-attention regions. We performed the task of replacing the 10bp motif centered on the selected region with “N” one motif at a time. We then calculated the difference from the unmasked sequence and considered the maximum absolute value of these differences as the impact of the masked motif.

Figure 9 shows the violin plot of the impact of the masked motifs between the high attention regions and low attention regions. The median difference in high-attention regions was 0.044, whereas in low-attention regions it was 0.005, indicating a clear separation in distributions. Statistical analysis using the Mann–Whitney U test confirmed a significant difference between the two groups (*p <* 0.001). The effect size was large, with a Cliff’s delta was 0.838, further supporting a substantial shift in the distribution of prediction differences between high- and low-attention regions. Among the 3,662 motifs with high attention, 461 motifs exhibited changes exceeding 0.2, a commonly used threshold indicating splicing alterations. In contrast, only 14 motifs with low attention showed such changes. This indicates that motifs with high attention are more likely to influence splicing than those with low attention.

**Fig. 9:**
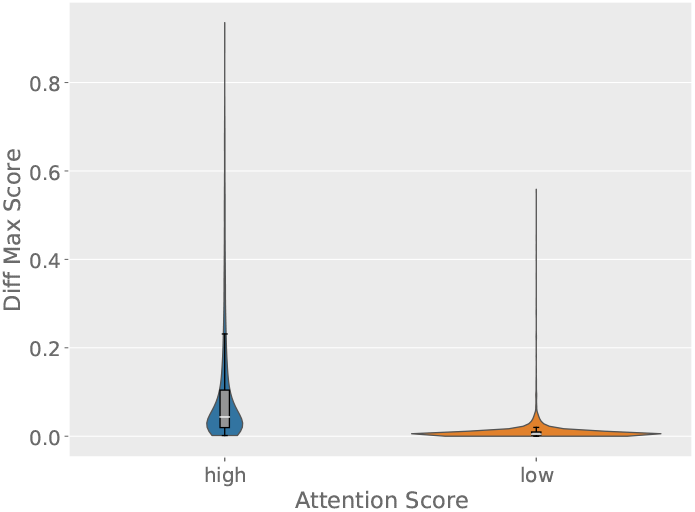
Prediction differences by high attention regions and low attention regions. The probability density is shown by violin plots, while internal box plots highlight the interquartile range and median (white line).

Therefore, we constructed a binary classification task to predict whether masking the corresponding region using the attention Z-score would yield a significant difference in the predicted values. The attention Z-score was calculated by multiplying the global attention value and the local attention value for each base, then using the mean and standard deviation of the product over the effective length excluding padding.

The AUROC obtained at multiple thresholds (0.05, 0.1, 0.2) ranged from 0.766 to 0.812, demonstrating that attention scores provide robust, threshold-independent rankings based on the magnitude of prediction differences (Supplementary Figure S8).

This suggests that attention captures monotonic information related to prediction sensitivity rather than reflecting threshold-specific artifacts.

Furthermore, for the high attention motifs used in this experiment, we searched for sequence motifs among those showing significant changes (0.1 or greater). For 10-bp motifs, we performed gap-free pairwise alignments for all possible pairs, created a graph by connecting motifs with scores above a threshold, and then clustered them using Markov Cluster Algorithm [41]. For each cluster, the motif with the highest degree was designated as the central motif, and motifs directly connected to it by edges were selected as representative motifs. Clusters were ranked based on the maximum number of bits achieved when creating logos for their representative motifs. For the top 5 clusters, the 10-bp motifs were extended by 10 bp on each side, and logos were created using the resulting 30-bp sequences. Using TOMTOM [42], we searched the JASPAR SPLICE (2020) [43] and Ray2013_rbp_Homo_sapiens [44] databases.

Two clusters showed similarity to RBP binding motifs, while the remaining three showed similarity to acceptor site motifs (Figure 10). Motif 1 showed similarity to binding motifs of RNA-binding proteins such as TIA1, U2AF2, and HuR. TIA1 primarily binds to the 3^′^ UTR but is also known to bind near intron donor sites, promoting splicing [45, 46, 47]. U2AF2 binds to the polypyrimidine tract (Py-tract) upstream of the 3^′^ splice site and is an essential factor for splicing initiation [48]. HuR contributes to mRNA stabilization and has also been reported to be involved in regulating alternative splicing in specific genes [47, 49, 50]. Motif 4, on the other hand, was similar to binding motifs for PABPC1, PABPC5, SART3, and others. PABPC5 and PABPC1 are proteins with high affinity for Poly-A sequences, and it has been suggested that these PABP family members are also involved in regulating alternative splicing within the nucleus [51]. Particularly under specific conditions, such as in cancer cells, these factors may contribute to changes in splicing patterns [52]. Furthermore, SART3 is an essential factor for the recycling of U4/U6 snRNPs, which constitute the spliceosome [53]. The recognition of these splicing-related factors via the motif indicates that the model captures the fundamental aspects of the splicing reaction. For motifs 2, 3, and 5, similarity to the acceptor site was detected. While the consensus sequence for the donor site is relatively short and highly conserved, the acceptor site’s “context” is defined by a broader and more diverse sequence information, including the upstream branch point and polypyrimidine tract. Therefore, this model is thought to have directed relatively higher attention toward the sequence features surrounding the acceptor site, which require more complex pattern recognition compared to the donor site. Furthermore, the detection of motifs associated with the acceptor site as multiple clusters suggests that the model may have individually learned the distinct sequence elements contributing to acceptor recognition and the subtle features of alternative splicing sites competing with constitutive splice sites.

**Fig. 10:**
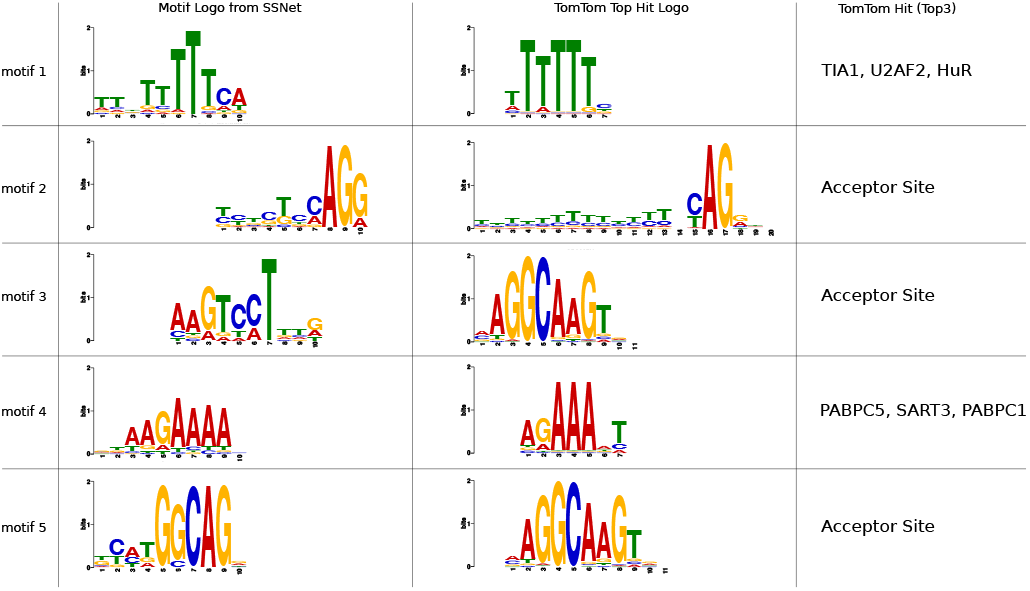
Top5 motif logo detected from high attention motifs clustering and TOMTOM top hit logo. TOMTOM top3 hits are also shown.

These results support the qualitative case studies presented in the previous section, indicating that attention highlights functionally influential sequence regions, although it should not be interpreted as a direct explanation.

### Long-Range Modeling in DMD Simulation and ClinVar Dataset

In this section, we comprehensively evaluate the ability of the proposed model’s hierarchical Transformer architecture to capture long-range genomic dependencies. While state-of-the-art (SoTA) CNN-based models like SpliceAI and Pangolin can theoretically process contexts up to 5 kb on either side of a target site, our framework expands this theoretical capacity up to 100 kb. To demonstrate the practical superiority of our architecture over deeply dilated convolutions, we performed two complementary experiments: a long-range decoy simulation utilizing the DMD gene, and a validation using real-world pathogenic mutations from the ClinVar database.

First, we conducted a long-range decoy experiment using the DMD gene. Deep intronic variants at ultra-long distances beyond the reach of SoTA models are extremely rare, and many involve variants creating pseudo-exons. This makes it difficult to directly assess the effectiveness of using ultra-long distances as input. Therefore, we focused on the exceptionally long intron 44 of the DMD gene and by introducing sequences that create decoy donor sites within the intron, we verified how far the model can capture variant effects. Specifically, we examined how much the probability of the actual donor site decreases due to the introduction of decoy donor sites.

The introduced sequence is a 198 bp decoy donor sequence combining a poly-A sequence (30 bp) + the sequence of exon 44 with the first 30 bp and last 3 bp removed + the consensus donor sequence “CAGGTAAGT” + the first 7 bp to 40 bp of intron 44. Regarding the exon 44 sequence, the ‘AG’ acceptor site capable of activation was replaced with ‘CG’ to prevent pseudo-exon formation. For the insertion position, distances from the true donor site of the decoy donor sequence were set in 100-base increments from 200 bp to 10 kb. Furthermore, for each distance, shifts were performed in 10-bp increments from -40 bp to 40 bp, creating nine mutated sequences for each distance to perform predictions.

Figure 11 shows the decrease in the probability of being a true donor site when decoy sequences are introduced at each distance for the SpliceAI, Pangoin, and SSNet_gtex_pangolin models. First, overall, the proposed model shows significant changes in predicted values at all distances, indicating it best captures the influence of decoy donor sequences. SpliceAI and Pangolin only show large changes when the decoy donor is in close proximity (200 bp). Beyond that, even within the receptive field range (<5 kb), they barely capture the effect of the decoy donor site on the constitutive donor site. Furthermore, since detecting changes beyond 5 kb is theoretically impossible, this value becomes 0.0. In contrast, the proposed model can continue to capture changes without significant attenuation even at ultra-long distances like 5 kb to 10 kb.

**Fig. 11:**
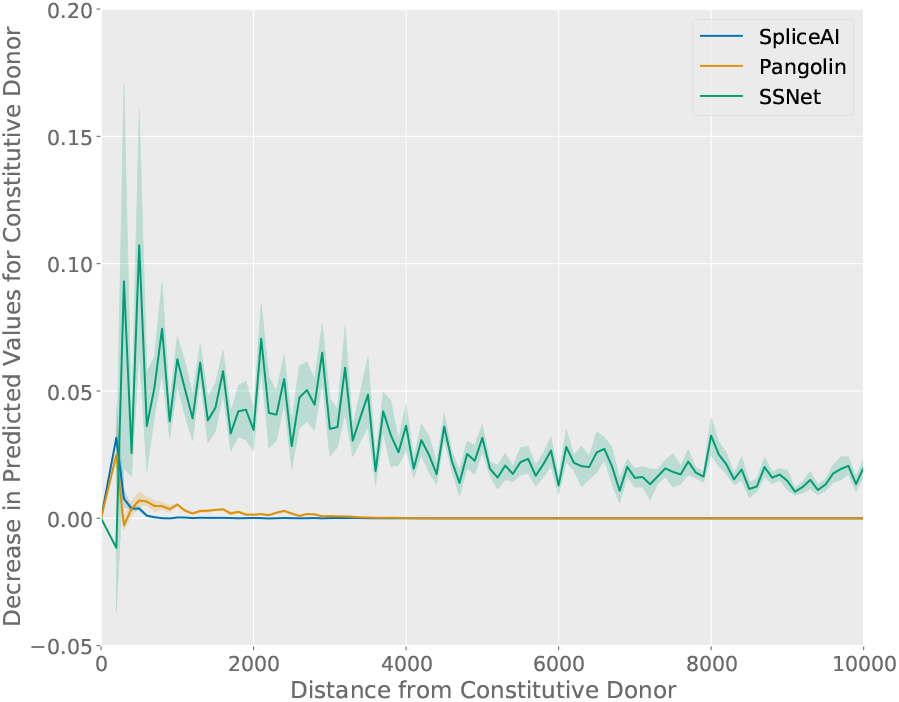
Changes in predicted values for true donor sites when decoy donor sequences are introduced at various distances. The predicted values for various shifts at each distance are shown as light-colored bands, and their average values are plotted as thick lines.

This simulation demonstrates that the proposed model captures the effects of variants at ultra-long distances beyond the receptive field of SoTA models. Furthermore, even within the receptive field, CNN-based SoTA models only capture effects up to a few hundred base pairs, revealing that the Transformer-based proposed model excels in considering long-range effects. This aligns with findings from theoretical studies in image processing tasks [54], which show that the proportion of the effective receptive field (ERF) within the theoretical receptive field is small for deep CNNs. While models like SpliceAI and Pangolin expand the effective receptive field through dilation compared to simple deep CNNs, this expansion is insufficient for capturing long-range effects. A model like the proposed one, utilizing a Transformer architecture, is necessary to more directly capture long-range information.

Next, to determine whether the architectural advantages observed in the DMD simulation translate to real-world genetic data, we benchmarked the models using real-world pathogenic variants extracted from the ClinVar database [55]. We analyzed variants annotated as “Pathogenic” or “Likely pathogenic” in the ClinVar dataset. For each variant, the distance to the nearest splice site was calculated, and only those located ≥ 1 kb away were retained. We then compared model prediction values between reference sequences and alternative sequences.

Recognizing that ClinVar contains diverse pathogenic mechanisms (e.g., missense or nonsense mutations) unrelated to splicing, evaluating all variants naively would dilute the underlying splicing signal. Furthermore, long-range pathogenic variants predominantly cause splicing defects through the activation of cryptic splice sites rather than a direct disruption of the nearest constitutive junction. To address these factors and isolate genuine splicing-disrupting mutations, we monitored the maximum alteration score (Max ΔScore) across the entire processed sequence window rather than restricting our focus to the closest constitutive splice site. To eliminate non-splicing artifacts without introducing algorithmic bias, we selected variants where either SpliceAI, Pangolin, or SSNet predicted a functional splicing disruption (Max ΔScore ≥ 0.05). To clearly visualize the distance-dependent sensitivity decay, the resulting profile was partitioned into two fine-grained distance windows (Figure 12).

**Fig. 12:**
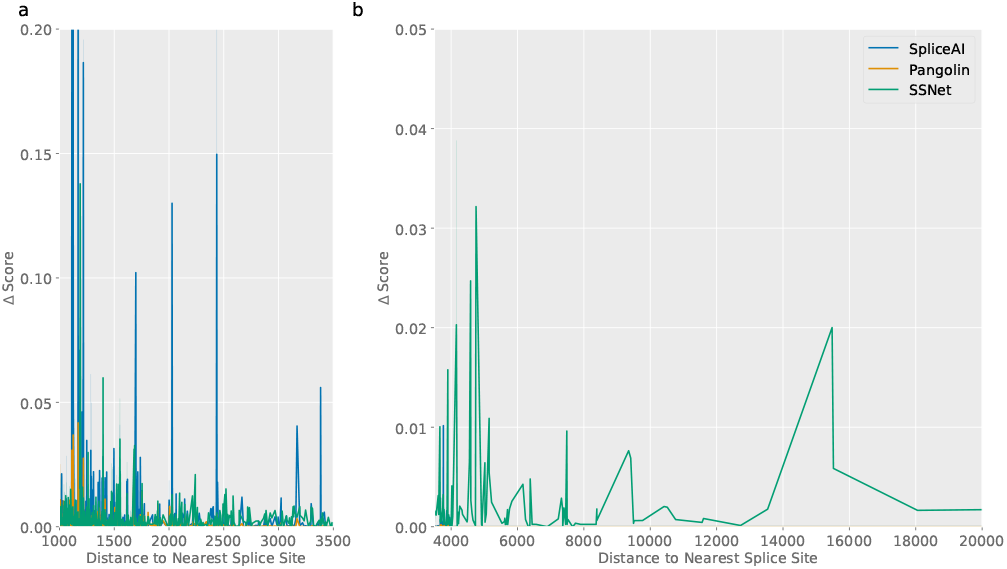
Distance-dependent decay of predicted splicing alterations (Max ΔScore) for pathogenic ClinVar variants. (a) In the proximal-to-intermediate range (1k–3.5k bp, scaled 0.0–0.2), all models demonstrate robust baseline predictions. (b) In the long-to-ultra-long range (3.5k–20k bp, scaled 0.0–0.05), SpliceAI and Pangolin sensitivity prematurely collapses to zero, while SSNet captures distinct functional signals.

The empirical distance profile unveils distinct architectural characteristics across three key genomic zones:

- **Zone I (1.0 kb – 3.5 kb):** The predictions of Pangolin and the proposed model show highly similar trends. Conversely, the sporadic, high-magnitude spikes observed exclusively in SpliceAI within this region are thought to reflect differences in training data definitions. SpliceAI is trained on datasets where any position utilized as a splice site in GTEx is defined as a positive instance, making its predictions extremely sensitive to mutational changes. In contrast, Pangolin and the proposed model learn splice site strengths as continuous values. Consequently, they tend to estimate mutational effects in a more gradual and quantitative manner, resulting in highly congruent behavior between these two models in the proximal range.
- **Zone II (3.5 kb – 5.0 kb):**Crucially, a striking divergence occurs in this window. Although this region resides strictly within SpliceAI and Pangolin’s theoretical ± 5 kb receptive field, its predicted values abruptly collapse to near-zero. This provides strong real-world validation for our simulation hypotheses. Specifically, it empirically demonstrates that for deep CNNs, the ERF is substantially narrower than the theoretical window. As a result, the variant positions fall outside the ERF well before reaching the throretical receptive field, leading to a premature loss of sensitivity. In contrast, our proposed model entirely circumvents this early collapse, maintaining stable sensitivity.
- **Zone III (5.0 kb – 20.0 kb):**Beyond the 5 kb theoretical limit where deep CNN models are structurally blind, the proposed model stands alone in its capacity to interpret distal regulatory variants. Most notably, it successfully captures a distinct signal originating nearly 15 kb away from the variant.

Taken together, these real-world validation results demonstrate that our hierarchical Transformer architecture not only broadens the theoretical window of splicing interpretation but effectively resolves the practical signal decay inherent to conventional deeply dilated convolutions, unlocking the ability to accurately interpret deep pathogenic mutations.

### Ablation Study

We conducted an ablation study to investigate how SSNet’s learning process contributes to this high performance and compared the performance on Gencode dataset (Table 3).

**Table 3.** Performance comparison on the Gencode test dataset for various SSNet models. SSNet_base denotes the model trained only on the Gencode dataset; SSNet_da is trained without intron/exon labels; SSNet_a0 is trained without applying class weighting to handle label imbalance; SSNet_g0 is trained without focal loss; SSNet_1k and SSNet_10k indicate models trained with input lengths of 1k and 10k nucleotides; SSNet_noconv, SSNet_nolocal, and SSNet_noglobal represent models with the convolutional, local attention, or global attention components removed, respectively. Values are reported as mean (95% confidence interval), estimated using gene-level bootstrap resampling.

| Model | Recall | Precision | F1 | Top-k | auPRC |
| --- | --- | --- | --- | --- | --- |
| SSNet_base | 0.937 (0.932-0.941) | 0.934 (0.929-0.938) | **0.935 (0.931-0.939)** | **0.940 (0.936-0.944)** | 0.943 (0.935-0.950) |
| SSNet_da | 0.920 (0.916-0.925) | 0.779 (0.771-0.787) | 0.844 (0.838-0.849) | 0.878 (0.873-0.883) | 0.904 (0.895-0.911) |
| SSNet_a0 | 0.919 (0.914-0.925) | 0.929 (0.924-0.934) | 0.924 (0.920-0.928) | 0.929 (0.924-0.933) | 0.937 (0.929-0.944) |
| SSNet_g0 | 0.930 (0.925-0.935) | **0.935 (0.930-0.939)** | 0.932 (0.928-0.936) | 0.937 (0.932-0.941) | 0.938 (0.930-0.945) |
| SSNet_1k | 0.804 (0.796-0.812) | 0.914 (0.908-0.921) | 0.856 (0.850-0.861) | 0.870 (0.865-0.875) | 0.887 (0.878-0.895) |
| SSNet_10k | 0.908 (0.902-0.914) | 0.930 (0.925-0.934) | 0.919 (0.914-0.923) | 0.925 (0.921-0.929) | 0.931 (0.923-0.938) |
| SSNet_noconv | 0.288 (0.280-0.297) | 0.617 (0.605-0.629) | 0.392 (0.384-0.402) | 0.460 (0.453-0.468) | 0.478 (0.469-0.487) |
| SSNet_noconv(opt) | 0.761 (0.753-0.768) | 0.691 (0.682-0.700) | 0.723 (0.717-0.729) | 0.746 (0.740-0.752) | 0.790 (0.782-0.799) |
| SSNet_nolocal | 0.890 (0.885-0.895) | 0.797 (0.789-0.804) | 0.841 (0.836-0.846) | 0.861 (0.857-0.866) | 0.887 (0.879-0.896) |
| SSNet_noglobal | 0.714 (0.704-0.724) | 0.901 (0.892-0.909) | 0.796 (0.789-0.804) | 0.835 (0.829-0.841) | 0.858 (0.849-0.866) |

First, we compared the SSNet training without intron/exon labels (SSNet_da). In this case, there was a significant performance drop in all metrics, with the largest drop in Precision. When training a language model as large as 100 kb, acceptor/donor labels alone are not sufficient information, and the addition of intron/exon labels may have improved performance by allowing learning of sequence features within the exon. In addition, the decrease in Precision was much larger than the decrease in Recall. This indicates that the intron/exon labels are helping to reduce false positives, while the acceptor/donor labels can learn the features of highly conserved motifs such as the GT-AG rule, but cannot learn the differences between spliced sites and non-spliced GT, AG dinucleotides. The addition of intron/exon labels would facilitate learning the context of whether or not a site is used as a splice site.

Next, we compared the performance of alpha and gamma, which are innovations in the loss function, with that of the case in which they were not used. Except for precision in SSNet_g0, which did not use gamma, SSNet_base performed the best in all indices, indicating that the weighting in the loss function contributes to the model performance. The difference between SSNet_a0 and SSNet_base is larger, indicating that weighting that resolves label imbalance is important for training data sets with very large label bias, such as spliced sites.

Then, a comparison was made for different input lengths. When the input length was set to 1k and 10k, the performance was worse for all indicators compared to when the input length was 100k. The performance improvement was more pronounced when the input length increased from 1k to 10k compared to 10k to 100k. This may be because more introns are fully included in the input with the increase from 1k to 10k.

Finally, we performed a layer-wise ablation study to assess the functional contributions of different model components. The model was partitioned into three major components: convolutional layers, local attention layers, and global attention layers. Based on this division, we trained three modified models—SSNet_noconv, SSNet_nolocal, and SSNet_noglobal—each lacking the respective component. The architectural details of ablation models (SSNet_noconv, SSNet_nolocal, SSNet_noglobal) are provided in Supplementary Figure S9.

For SSNet_noconv, performance deteriorated substantially across all evaluation metrics. To partially mitigate this degradation, we introduced an additional variant, SSNet_noconv(opt), in which the learning rate was increased fivefold to 0.00025. Although this optimized variant still exhibited the lowest overall performance among all ablation models (except for Recall), it showed a marked improvement compared to the original SSNet_noconv without learning-rate adjustment. These results indicate that the convolutional layers, forming the initial part of the model, not only extract and learn local motifs, such as the GT–AG signal indispensable for splice sites, but also play a critical role in stabilizing and facilitating overall training. Without convolutional layers, learning fails to progress effectively at a small learning rate, leading to substantial performance degradation.

Comparing SSNet_nolocal and SSNet_noglobal reveals that the latter exhibits inferior performance, indicating that global attention, which captures long-range dependencies, contributes more substantially to overall performance than local attention. As convolutional layers already capture a certain degree of local motif information, the local attention layers appear to serve a more auxiliary role. The ablation results further reveal distinct effects: SSNet_nolocal exhibits a marked decrease in Precision, whereas SSNet_noglobal exhibits a pronounced decline in Recall. These findings highlight the complementary roles of local and global attention. Local attention enables the model to capture fine-grained contextual dependencies around splice sites; without it, the model may overlook relatively weakly conserved motifs beyond the canonical GT–AG rule, increasing False Positives. Conversely, global attention enables the model to learn long-range dependencies and broader contextual relationships. Its absence hampers detection of splice sites that are locally weak but supported by long-range contexts, thereby increasing False Negatives.

In summary, this layer-wise ablation study confirms the functional contributions of each model component. The findings demonstrate that both local feature extraction and the modeling of long-range dependencies via global attention are essential for accurate splice site prediction.

### Inference Time and Memory Comparison

To assess the computational efficiency of the proposed model during inference, we measured and compared inference times for SSNet, SpliceAI, Spliceformer, and SpliceTransformer. Pangolin was excluded, as its architecture is nearly identical to SpliceAI except for the final classification layer. SpliceBERT is capable of processing only very short input lengths. It could only run without OOM on the shortest input length in this experiment (10k), so it was excluded from the figure, and only the values for 10k are shown in the text. For a fair comparison, all measurements were conducted on an NVIDIA H100 GPU with CUDA 11.8, both with and without mixed precision. As the main component of SpliceTransformer, the Sinkhorn Transformer, does not support mixed precision, its measurements were obtained only without mixed precision.

Figure 13 a illustrates inference times across various input lengths. SpliceBERT shows slowest inference time (0.8991 seconds for 10kb without mixed precision and 0.5164 seconds with mixed precision). SpliceTransformer exhibits extremely long inference times for all input lengths due to its Transformer-based architecture, which attends to sequences ranging from thousands to tens of thousands of positions via the attention mechanism. Spliceformer, also Transformer-based, runs faster than SpliceTransformer because its Attention mechanism considers only the top 512 splice site candidates, independent of sequence length. Under default mixed precision, SSNet is the second fastest after SpliceAI. Despite processing the entire input sequence, SSNet’s hierarchical architecture reduces the effective input length for attention computation, enabling efficient inference. Notably, SSNet evaluates the full input sequence, giving it a receptive field equal to the input length, whereas SpliceAI’s receptive field is fixed at 10k.

**Fig. 13:**
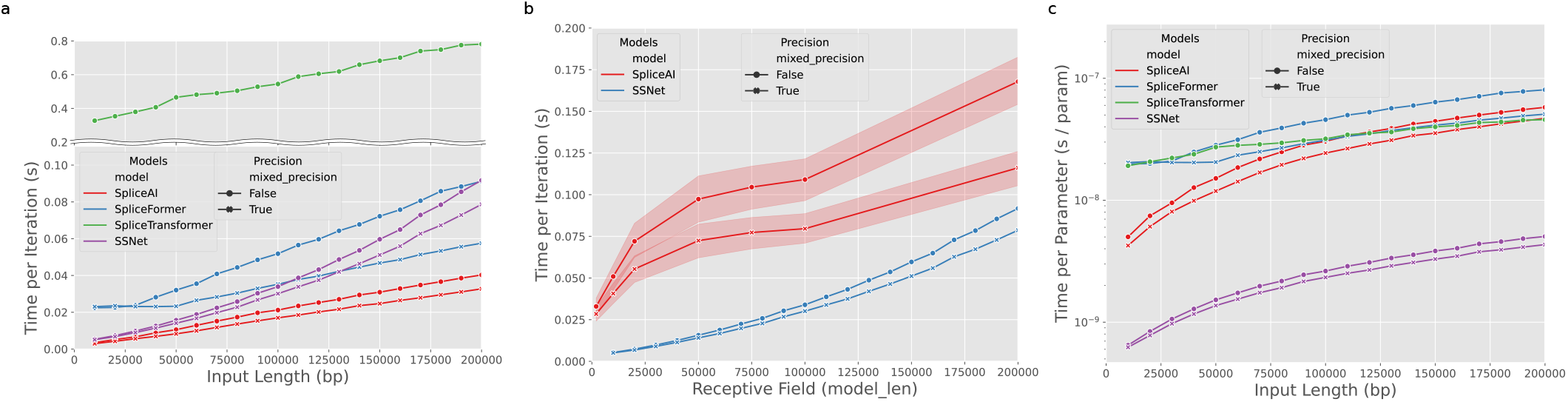
(a) Inference time of different models (SpliceAI, Spliceformer, SpliceTransformer, SSNet) across various input sequence length. (b) Inference time versus receptive field size for SSNet and SpliceAI. (c) Inference time normalized by model parameters across input lengths.

We next investigated how inference times change across various receptive fields for the two fastest models in the previous comparison, SpliceAI and SSNet (Figure 13 b). For SpliceAI, we extended the official 2k and 10k architectures by adding convolutional layers to achieve receptive fields of 20k, 50k, 75k, 100k, and 200k. Inference times at various input lengths for each SpliceAI variant are shown in light blue widths. While SpliceAI was faster in inference times across various input lengths, SSNet consistently showed faster inference times across various receptive fields. This indicated the computational efficiency of the proposed model.

Finally, to evaluate the relationship between model size (number of parameters) and inference time, we plotted inference time divided by the number of parameters across different input lengths (Figure 13 c). SpliceBERT shows almost the same performance with SpliceTransformer and Spliceformer (3.74×10^−8^ for 10kb without mixed precision and 2.15 × 10^−8^ with mixed precision). Although SpliceTransformer had the longest absolute inference times, normalizing by its relatively large number of parameters brought its performance in line with Spliceformer and SpliceAI. SSNet demonstrated high scalability relative to model size, achieving exceptionally low inference time per parameter.

The comparison of GPU memory consumption relative to input length is provided in Supplementary Figure S10. Since SSNet employs the Transformer’s attention mechanism, it entails a practical trade-off involving higher absolute memory consumption compared to CNN-based models like SpliceAI. However, when utilizing a vanilla BERT (e.g., SpliceBERT), injecting long-range contexts exceeding 10 kb immediately triggers OOM errors due to the quadratic scaling of attention computation. In contrast, SSNet dramatically mitigates this memory explosion through its hierarchical Transformer architecture. Although its absolute memory footprint is larger than that of CNNs, SSNet achieves vastly superior memory efficiency compared to vanilla Transformers, successfully bounding long-range dependency learning within a practical resource scale manageable on standard hardware environments.

## Discussion

We introduced SpliceSelectNet, the first hierarchical Transformer model capable of handling long-range dependencies for splice site prediction. To our knowledge, hierarchical attention mechanisms have rarely been applied in bioinformatics tasks, and no prior study has explicitly adopted such a design for splice site prediction. Thus, our architecture represents a novel contribution by enabling dense attention over long genomic sequences while remaining computationally efficient.

The success of SSNet’s in three benchmark datasets can be attributed to several innovations. The first one is the usage of intron/exon label in the first step of training. In large DNA language models, having the model perform the subtasks associated with donor/acceptor site predictions simultaneously increases the generality of the model and reduces overfitting. It also made it possible to capture sequence features within the exon, and the model was able to learn the context in which the splice site was present. This improved the model’s ability to distinguish between a real splice site and just a GT, AG dinucleotide, and reduced false positives. The second one is loss function design. The use of balanced cross-entropy and focal loss addressed the class imbalance issue inherent in splicing datasets. Since we often face the class imbalance problem in the field of bioinformatics, this technique can be applied to other biological prediction tasks. The third one is data augmentation. The incorporation of GTEx and Pangolin datasets enhanced the model’s ability to predict both constitutive and alternative splice sites. Based on our evaluation, we provide a practical recommendation for users applying SSNet to their research. We recommend using the SSNet_gtex_pangolin model as the default choice. Our results indicate that this configuration offers the most robust generalization performance, effectively capturing both the alternative splicing sites provided by the GTEx dataset and the splice site usage information derived from the Pangolin dataset. The SSNet_pangolin_gtex model remains a valid alternative for tasks strictly focusing on detection of the generation of new splice sites with high sensitivitys. The fourth one is longer input lengths. By extending the input length to 100 kb, the model effectively captured long-range dependencies. The final one is the design of model architecture. Our model consists of convolutional layers, local attention layers and global attention layers. Layer-wise ablation study showed that each component has its own important role and not only local feature extraction but also global context greatly contributed to highly accurate splice site predictions. An ablation study showed that each of these factors contributed significantly to the overall performance, highlighting their importance in model development.

SSNet’s hierarchical attention mechanism proved essential for handling long-range dependence, an important aspect of splicing control. Comparisons with SpliceAI and Pangolin in in-silico mutagenesis of DMD gene revealed that SSNet’s Transformer architecture allows more context information and enables better performance for mutations involved in complex splicing control mechanisms and for long-distance mutations. SSNet is also, to our knowledge, the first attempt in bioinformatics to use a hierarchical attention mechanism. This structure solves the limitation to the input length, which is a problem when dealing with DNA sequences while keeping the computational complexity low. Although the number of parameters is much larger than those of other existing models, the introduction of the hierarchical attention mechanism allows inference at almost the same or faster speed.

Furthermore, the attention heatmaps of SSNet paves the way for understanding splicing mechanisms. For example, in BRCA1 Exon 10, attention weights revealed a potential regulatory region upstream of the cryptic splice site, providing new insight into splicing activity by mutation. Since the model uses dense attention weights, visualization of the attention are intuitive, making the model easy to understand for experts in the fields of biology and medicine. The advantage is that the attention weights are the result of intermediate calculations during inference, so there is no need to perform additional calculations for interpretation. In addition, attention maps can also be leveraged to explore sequence motifs that are important for splicing regulation. By identifying which regions of the sequence receive higher attention weights, we can uncover previously unknown motifs or enhance the understanding of known splicing regulatory elements. For variants of uncertain significance, attention allows us to identify how the mutation affects splicing regulation, contributing to the elucidation of disease mechanisms. Indeed, our case studies on the IgM and FAS genes demonstrated that SSNet accurately reproduced the effects of ESE and ISE mutations, in line with experimental observations. These results confirm that the model not only predicts splicing outcomes but also provides mechanistic insights into cis-regulatory elements, bridging computational prediction with biological interpretation.

Recently, the advent of genomic foundation models, such as Borzoi [56] and AlphaGenome [57], has enabled the modeling of ultra-long-range contexts exceeding 100 kb, achieving exceptionally high predictive performance. Backed by massive parameter scales and computing resources, these foundation models possess an overwhelming advantage in comprehensively mapping global genomic contexts and multi-layered epigenomic dynamics. However, deploying such giant models strictly demands high-end computational infrastructure, such as multi-GPU clusters. In contrast, our proposed SSNet adopts a relatively lightweight, hierarchical Transformer architecture specifically optimized for splice-site prediction. While maintaining high accuracy, SSNet can efficiently process long-range dependencies (up to 100 kb) with fast inference times on single-GPU setups. Furthermore, while giant foundation models often suffer from a “black box” limitation due to their extreme architectural complexity, SSNet provides high interpretability by allowing direct visualization of attention maps. This enables researchers to trace exactly which long-range nucleotide signals drive the model’s predictions. Combining localized practical deployment with clear biological explainability, SSNet offers a powerful, complementary alternative to resource-intensive genomic foundation models.

The architecture of SSNet is versatile and could be extended to other genomic tasks, such as the prediction of transcription factor binding sites, chromatin accessibility, and epigenomic modifications. In particular, since it can take input sequences as long as 100k, it is expected to perform well in prediction tasks where long-range interactions are important. One promising future direction is to enhance the prediction of alternative splicing events in a tissue-specific manner. By integrating tissue-specific RNA-seq datasets and splicing regulatory element annotations, SSNet could identify key elements that drive differential splicing across tissues. This could provide novel insights into the regulatory mechanisms underlying splicing and uncover potential therapeutic targets for diseases associated with aberrant splicing. Another future plan is to develop a general-purpose DNA language model using this architecture. A DNA large language model with practical computational speed could be applied to a variety of downstream tasks.

In this paper, our model outperforms the current SoTA in both splice site prediction and aberrant splicing prediction. With its dense attention mechanism, SpliceSelectNet is poised to be a valuable tool for researchers studying gene regulation and disease-associated splicing variants. Moreover, its architecture can be generalized to other DNA sequence tasks, opening the door to broader applications as a versatile DNA language model.

## Author contributions statement

Y.M. conducted the majority of the research, including conceptualization, methodology development, data analysis, and manuscript drafting. K.N. supervised the project, provided critical guidance throughout the research process, and contributed to the manuscript’s revision and finalization. Both authors reviewed and approved the final version of the manuscript.

## Acknowledgments

This work was supported by JST SPRING, Grant Number JPMJSP2108. We also acknowledge the Human Genome Center, the Institute of Medical Science, the University of Tokyo, for providing the super-computing resource (http://sc.hgc.jp/shirokane.html).

## Data Availability Statement

The raw data used for model training were obtained from publicly available datasets as described in previous studies (e.g., [22] for Gencode Dataset, [23, 24] for GTEx Dataset, [25] for Pangolin Dataset). For the Gencode and GTEx datasets, we utilized the preprocessed datasets provided by SpliceAI [9]. For the Pangolin dataset, we processed the original RNA-seq data before use.

All code for data pre-processing, model training, and inference, as well as prediction results for all datasets and codes for reproducing the results are publicly available at https://doi.org/10.5281/zenodo.20407617.

