## Supplemental Figures and Tables for "SpliceSelectNet: A Hierarchical Transformer-Based Deep Learning Model for Splice Site Prediction"

### Supplementary Figures

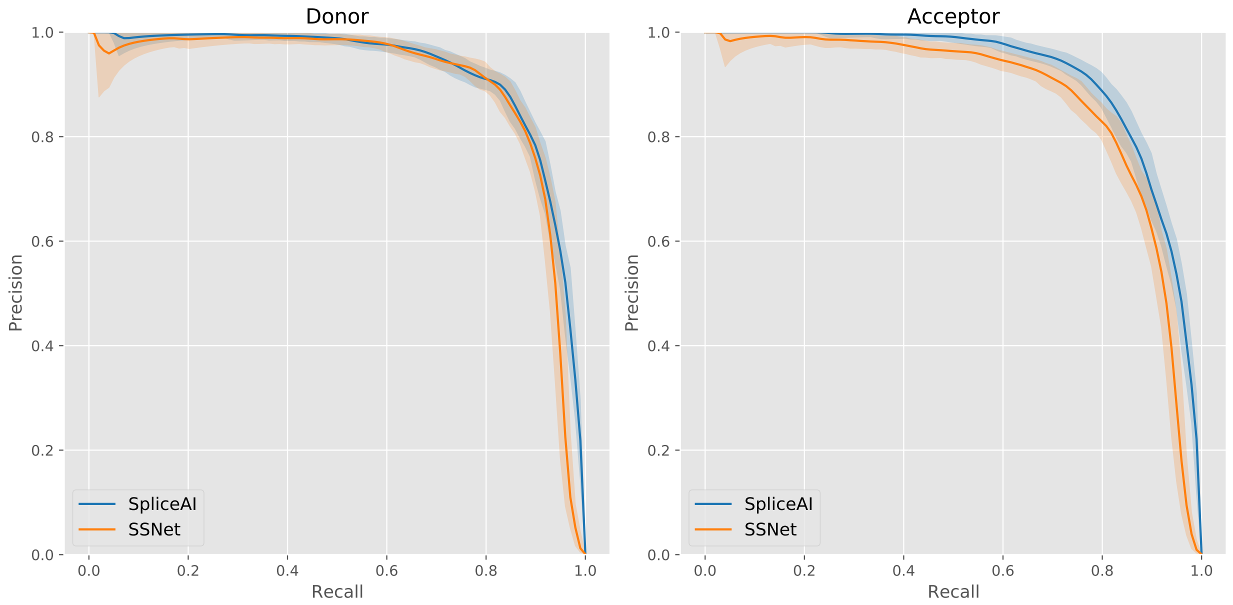


Supplementary Figure S1: Performance of SSNet_base and SpliceAI on lincRNA. PR curves are shown for donor and acceptor sites. Shaded regions indicate 95% confidence intervals estimated by stratified bootstrap resampling.

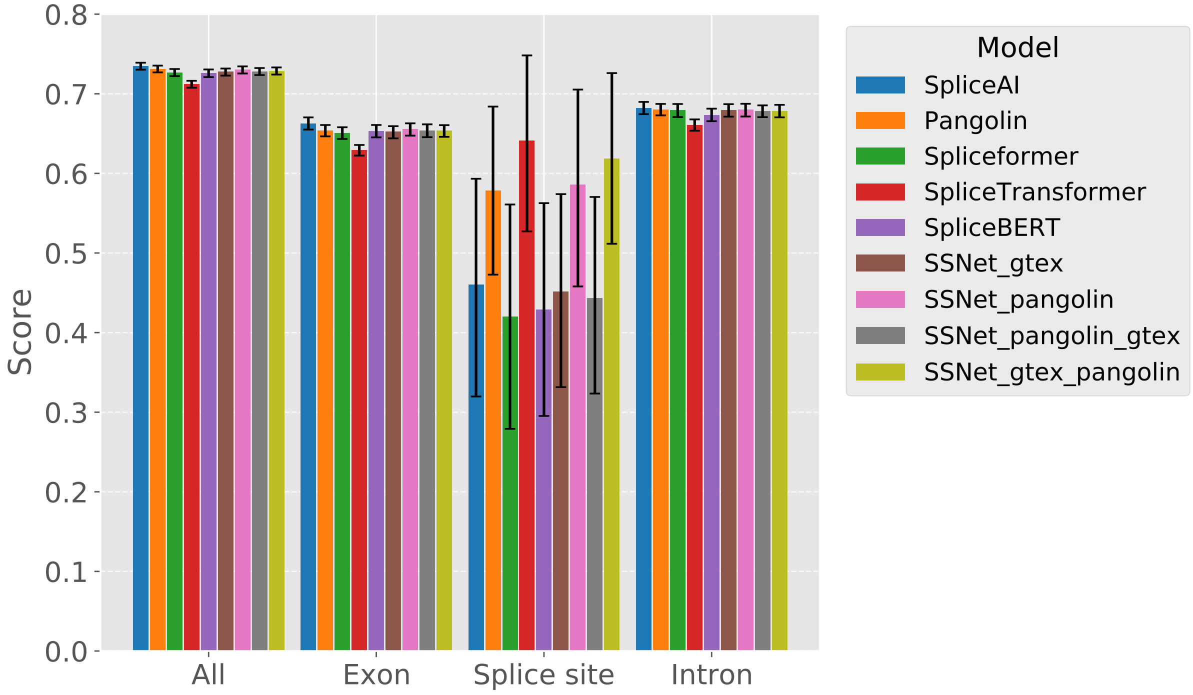


Supplementary Figure S2: Performance of all SSNet variants and state-of-the-art models on SpliceVarDB. AUROC values are shown for all variants combined and stratified by variant location (Exon, SpliceSite, Intron). SSNet variants include SSNet_gtex, SSNet_pangolin, SSNet_gtex_pangolin, and SSNet_pangolin_gtex. Other models include SpliceAI, Pangolin, Spliceformer, SpliceTransformer, and SpliceBERT. Error bars indicate 95% confidence intervals estimated by stratified bootstrap resampling.

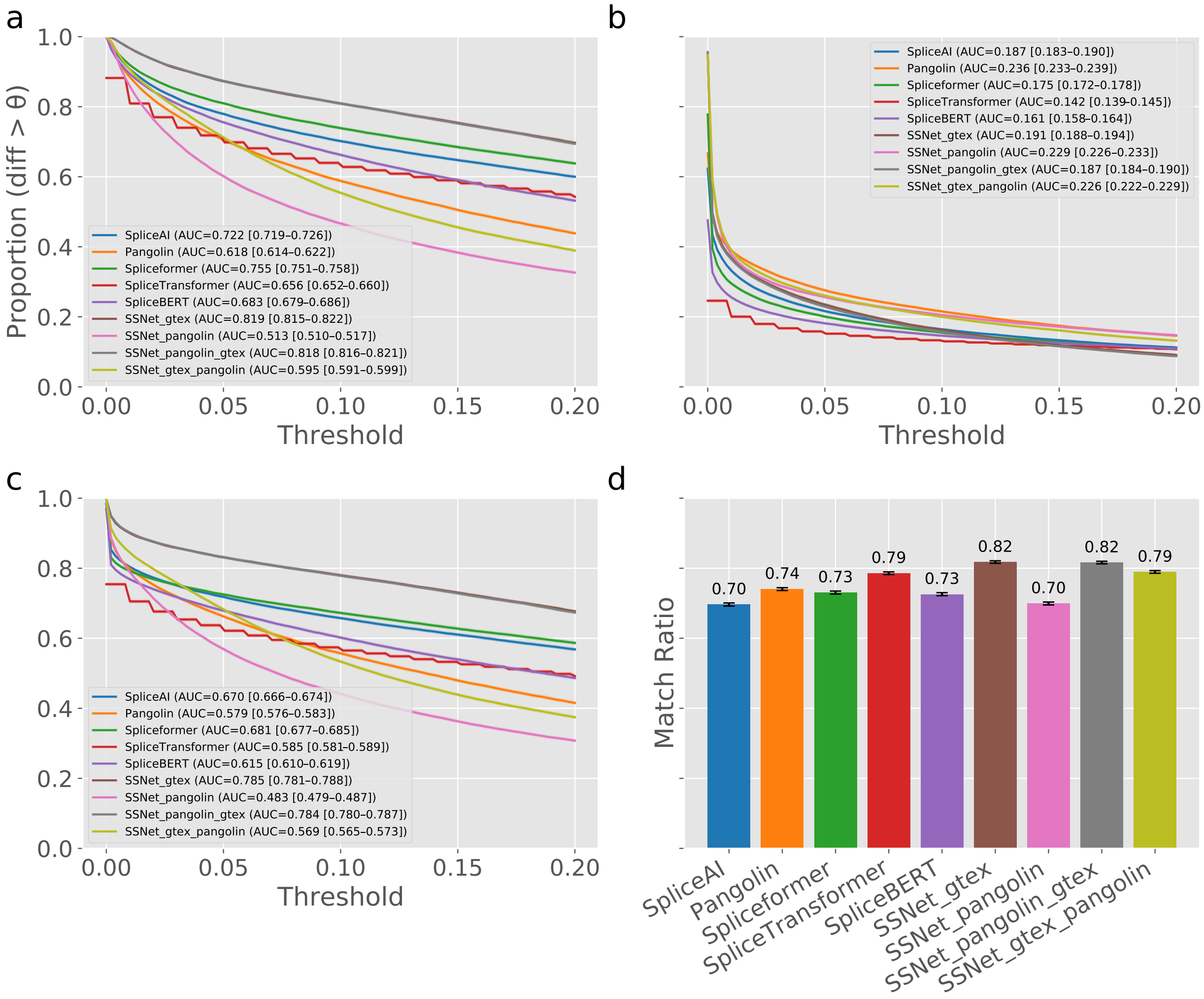


Supplementary Figure S3: Performance of all SSNet variants and state-of-the-art models on SSCVDB. Proportion of predicted splicing changes exceeding thresholds (0.0–0.2) is shown for (a) the entire sequence, (b) original splice sites (Hijacked_SS), and (c) newly created splice sites (Primary_SS) in SSCVDB. (d) The probability that the maximum predicted change occurs at Primary_SS. Error bars indicate 95% confidence intervals estimated by stratified bootstrap resampling.

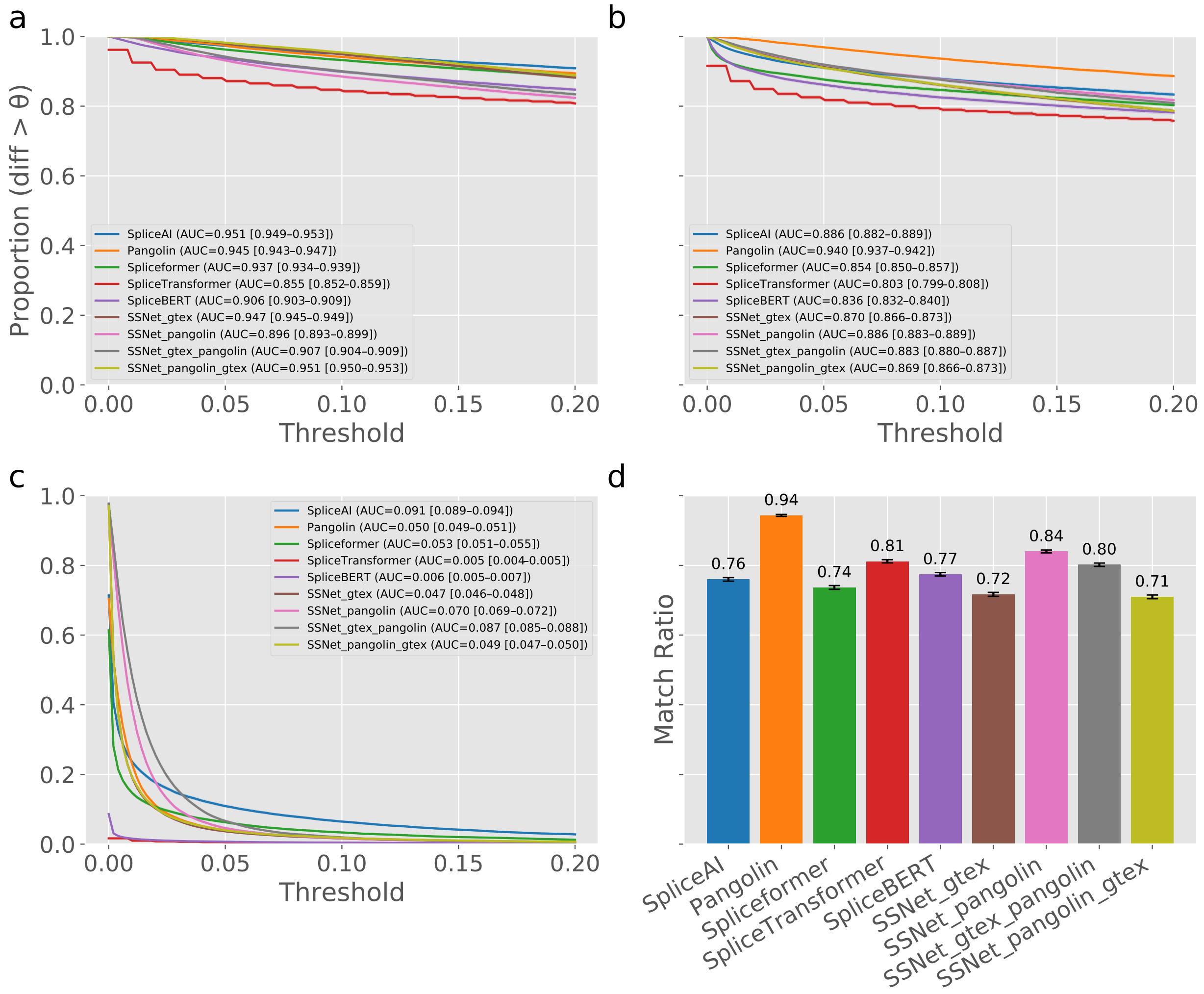


Supplementary Figure S4: Intron retention detection performance of all SSNet variants and state-of-the-art models on IRAVDB. Proportion of predicted splicing changes exceeding thresholds (0.0–0.2) is shown for (a) the entire sequence, (b) target splice sites (target_pos), and (c) their partner splice sites (partner_pos). (d) The probability that the maximum predicted change occurs at target_pos. Error bars indicate 95% confidence intervals estimated by stratified bootstrap resampling.

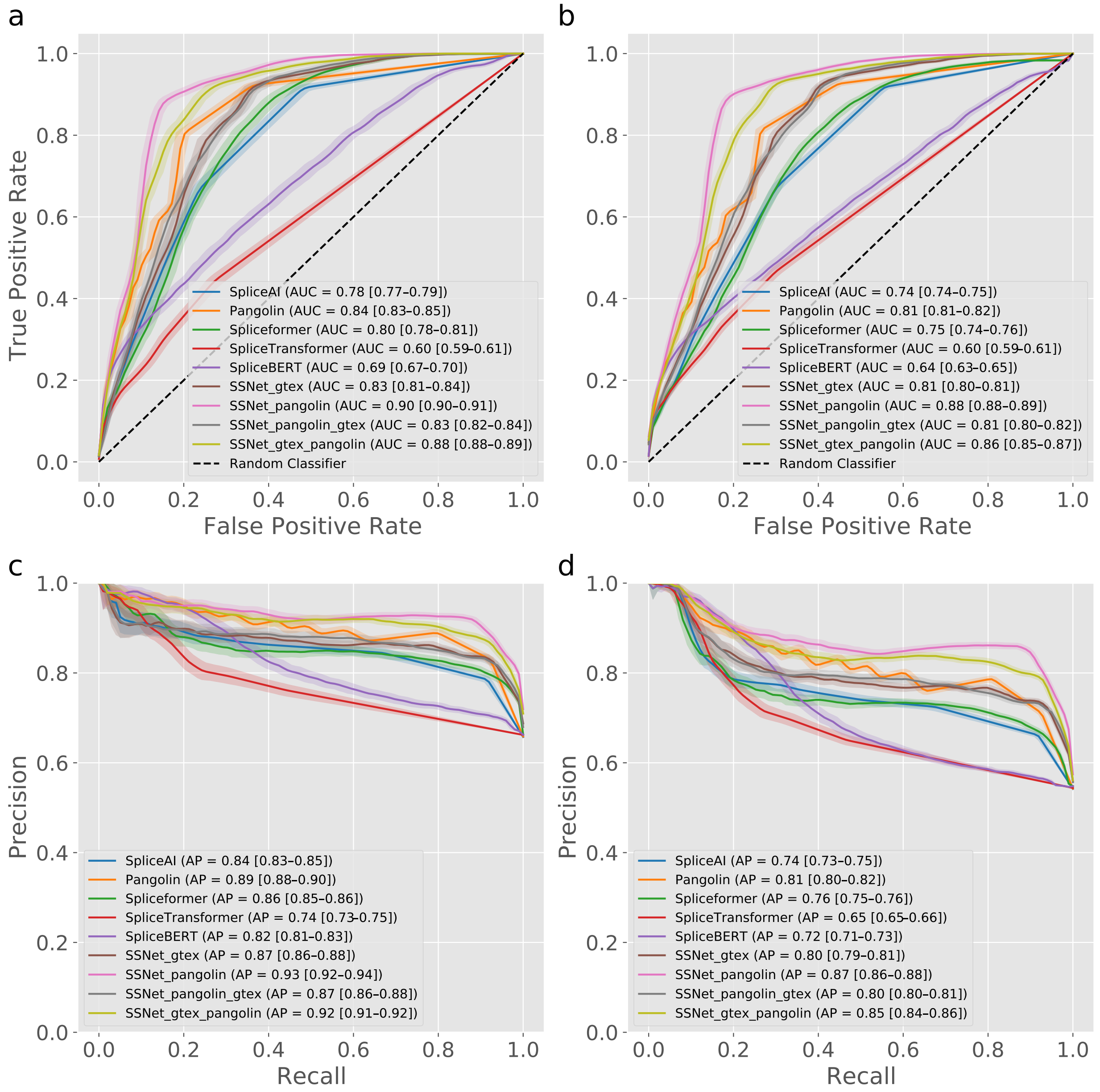


Supplementary Figure S5: Performance of all SSNet variants and state-of-the-art models on the BRCA dataset. Shaded regions indicate 95% confidence intervals estimated by stratified bootstrap resampling. (a) ROC curves for expert label. (b) ROC curves for all label. (c) PR curves for expert label. (d) PR curves for all label.

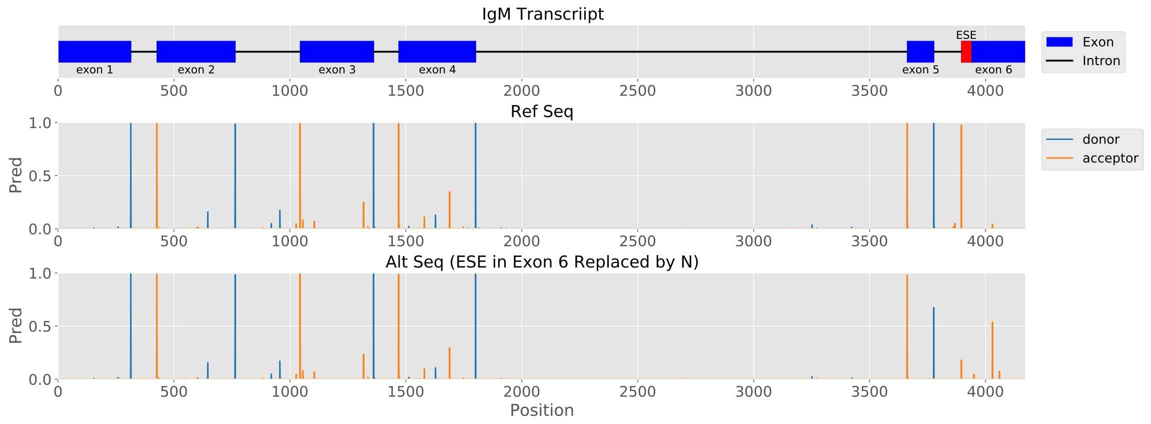


Supplementary Figure S6: SpliceAI predictions for the mouse IgM gene. Top: gene structure with introns (black lines) and exons (blue boxes); the 42 bp ESE at the 5' end of Exon 6 is highlighted in red. Bottom: predicted donor (blue) and acceptor (orange) probabilities for the wild-type sequence and with the ESE region masked with "N". Masking the ESE abolishes the acceptor site at the 5' end of Exon 6 (red box).

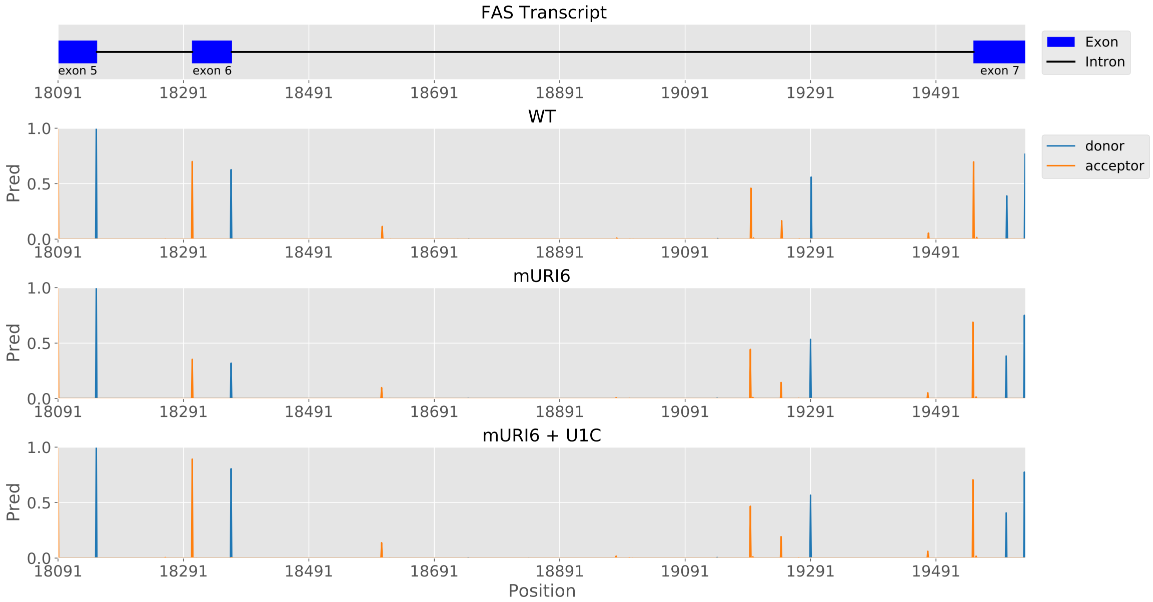


Supplementary Figure S7: SpliceAI predictions for FAS wild-type, mURI6, and mURI6+U1C sequences. The red boxes indicate regions where splice site probabilities are altered by mutations.

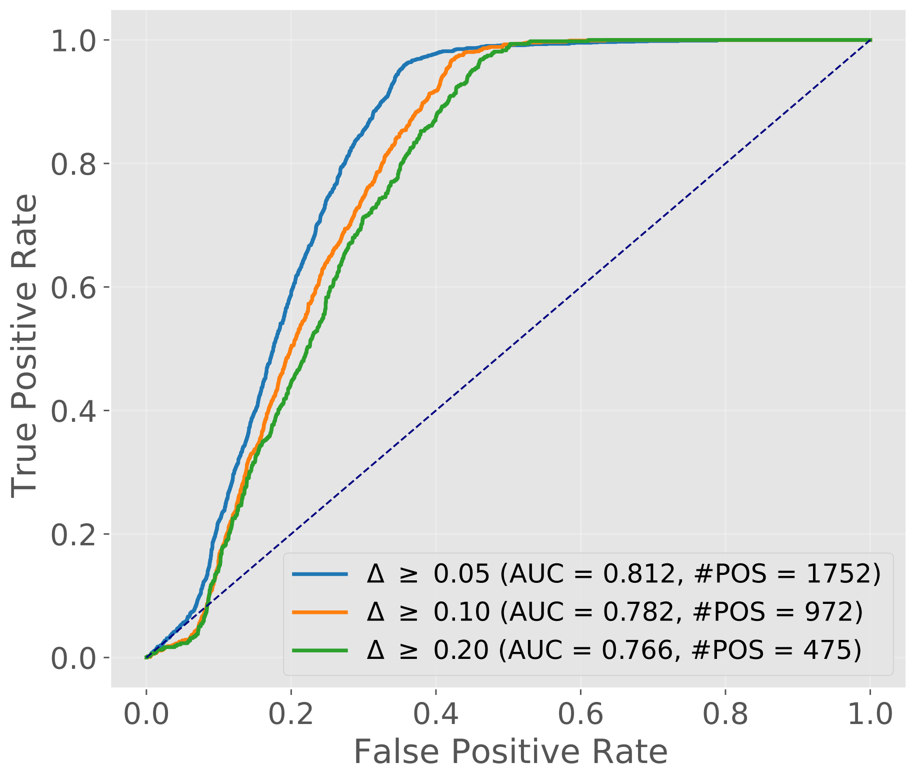


Supplementary Figure S8: ROC curves for prediction difference values based on attention z-score at multiple thresholds (0.05, 0.1, 0.2).

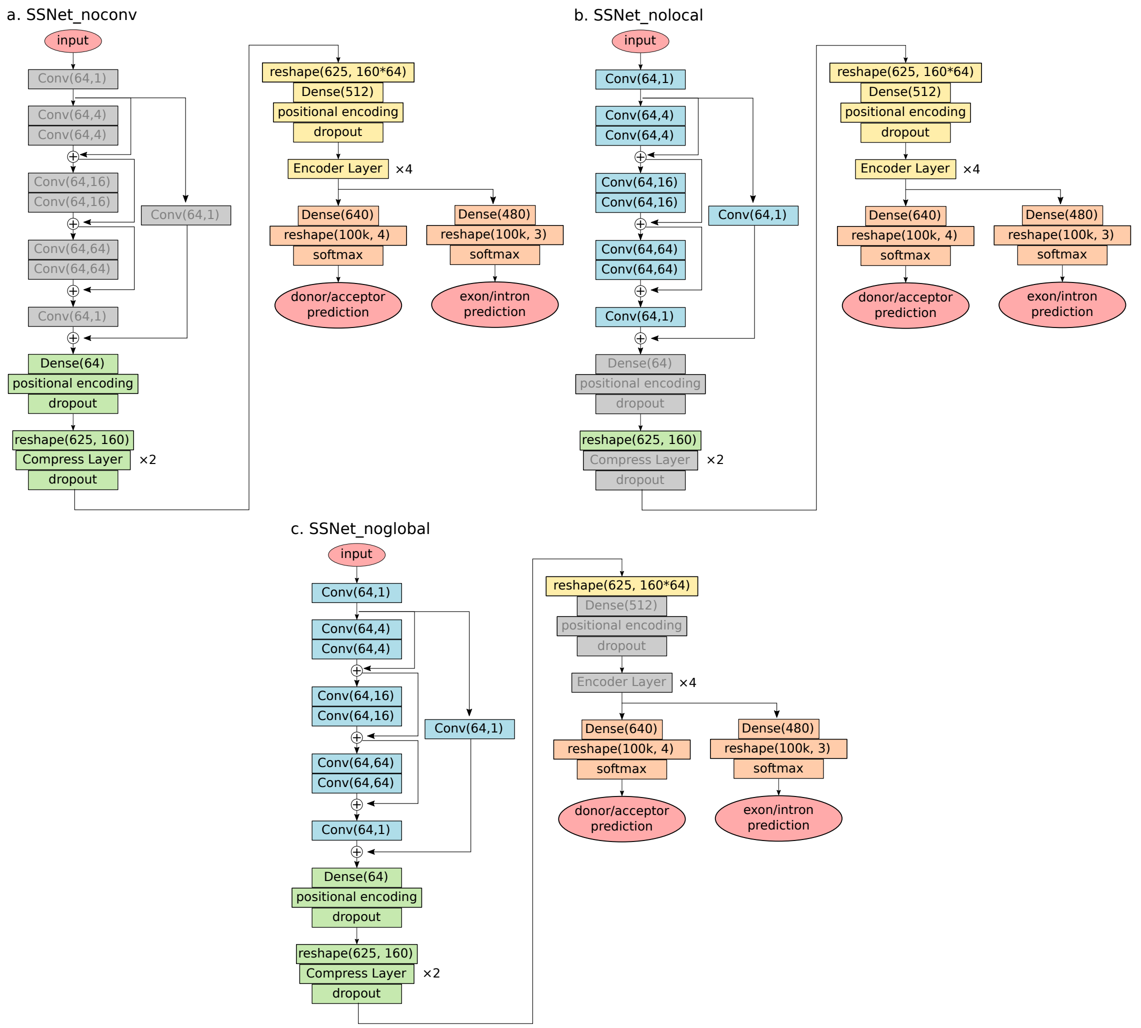


Supplementary Figure S9: Layer-wise ablation study of SSNet architecture. This figure highlights which components of the model (Convolutional layers, Local Attention, Global Attention) were used or commented out in the ablation experiments (SSNet_noconv, SSNet_nolocal, SSNet_noglobal). Performance metrics are reported in Table 1 of the main text.

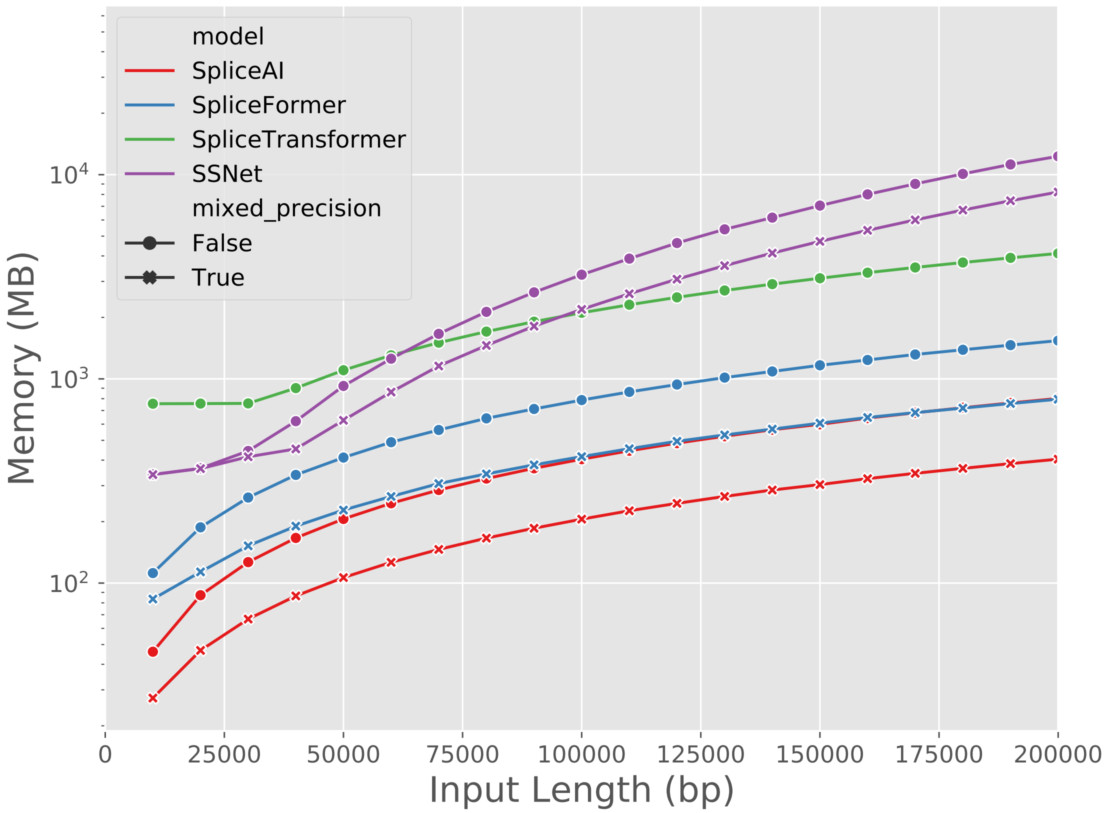


Supplementary Figure S10: Memory consumptions of different models (SpliceAI, Spliceformer, SpliceTransformer, SSNet) across various input sequence length.

### Supplementary Tables

Supplementary Table S1: Training Data Summary. Summary of datasets used for model training. Donor and acceptor counts are derived from labeled splice sites, and for Pangolin, the Average SSE (donor/acceptor) is reported.

| Dataset | #Genes | #Donor | #Acceptor | Ave SSE (donor) | Ave SSE (acceptor) |
|---|---|---|---|---|---|
| Gencode | 13384 | 130795 | 130795 | - | - |
| GTEx | 13385 | 193699 | 197804 | - | - |
| Pangolin | 13384 | 158621 | 156361 | 0.612 | 0.606 |

Supplementary Table S2: Validation Data Summary for SVDB. The number of genes and positive/negative variants for all regions and region-specific (Exon, SpliceSite, Intron) are shown. Label 3 shows splice-altering variants, Label 2 shows low frequency splice-altering ones, and Label 1 shows not splice-altering ones.

| Region | #Variants | #Genes | #label=3 | #label=2 | #label=1 |
|---|---|---|---|---|---|
| All | 48104 | 7538 | 12943 | 24282 | 10804 |
| Exon | 22859 | 4544 | 3357 | 13697 | 5740 |
| Splice Site | 7334 | 4350 | 5800 | 1511 | 22 |
| Intron | 17911 | 3990 | 3786 | 9074 | 5042 |

Supplementary Table S3: Validation Data Summary for BRCA. The number of genes and positive/negative variants for expert and all labels are shown.

| Label Type | # Variants | #Genes | #Pathogenic | #Benign | #Unknown |
|---|---|---|---|---|---|
| label_expert | 72467 | 2 | 4909 | 2544 | 65014 |
| label_all | 72467 | 2 | 7966 | 6619 | 57882 |

Supplementary Table S4: Validation Results for SVDB. TP, FP, TN, FN counts at fixed threshold are provided, along with AUROC and AUPRC. Results are grouped by region.

| Region | Model | #TP | #FP | #TN | #FN | AUROC | AUPRC |
|---|---|---|---|---|---|---|---|
| All | SpliceAI | 14765 | 259 | 10620 | 22460 | 0.735 | 0.916 |
| | Pangolin | 14629 | 210 | 10668 | 22592 | 0.731 | 0.918 |
| | Spliceformer | 14596 | 303 | 10576 | 22629 | 0.727 | 0.913 |
| | SpliceTransformer | 13156 | 178 | 10701 | 24069 | 0.712 | 0.912 |
| | SpliceBERT | 13942 | 242 | 10637 | 23283 | 0.726 | 0.912 |
| | SSNet_gtex | 14145 | 256 | 10623 | 23080 | 0.728 | 0.913 |
| | SSNet_pangolin | 12455 | 94 | 10785 | 24770 | 0.730 | 0.915 |
| | SSNet_gtex_pangolin | 13013 | 103 | 10776 | 24212 | 0.728 | 0.915 |
| | SSNet_pangolin_gtex | 14252 | 258 | 10621 | 22973 | 0.728 | 0.913 |
| Exon | SpliceAI | 3557 | 115 | 5690 | 13497 | 0.663 | 0.865 |
| | Pangolin | 3519 | 105 | 5700 | 13535 | 0.654 | 0.866 |
| | Spliceformer | 3490 | 137 | 5668 | 13564 | 0.651 | 0.860 |
| | SpliceTransformer | 2763 | 77 | 5728 | 14291 | 0.629 | 0.855 |
| | SpliceBERT | 3256 | 126 | 5679 | 13798 | 0.653 | 0.860 |
| | SSNet_gtex | 3244 | 131 | 5674 | 13810 | 0.652 | 0.859 |
| | SSNet_pangolin | 2292 | 37 | 5768 | 14762 | 0.655 | 0.863 |
| | SSNet_gtex_pangolin | 2689 | 40 | 5765 | 14365 | 0.653 | 0.862 |
| | SSNet_pangolin_gtex | 3304 | 134 | 5671 | 13750 | 0.653 | 0.860 |
| Splice Site | SpliceAI | 7272 | 21 | 2 | 39 | 0.459 | 0.996 |
| | Pangolin | 7265 | 21 | 2 | 46 | 0.576 | 0.998 |
| | Spliceformer | 7274 | 23 | 0 | 37 | 0.424 | 0.995 |
| | SpliceTransformer | 7223 | 22 | 1 | 88 | 0.640 | 0.998 |
| | SpliceBERT | 7251 | 23 | 0 | 60 | 0.429 | 0.996 |
| | SSNet_gtex | 7281 | 21 | 2 | 30 | 0.458 | 0.997 |
| | SSNet_pangolin | 7250 | 21 | 2 | 61 | 0.585 | 0.998 |
| | SSNet_gtex_pangolin | 7254 | 21 | 2 | 57 | 0.618 | 0.998 |
| | SSNet_pangolin_gtex | 7280 | 21 | 2 | 31 | 0.442 | 0.996 |
| Intron | SpliceAI | 3936 | 123 | 4928 | 8924 | 0.682 | 0.865 |
| | Pangolin | 3845 | 84 | 4966 | 9011 | 0.680 | 0.870 |
| | Spliceformer | 3832 | 143 | 4908 | 9028 | 0.679 | 0.863 |
| | SpliceTransformer | 3170 | 79 | 4972 | 9690 | 0.661 | 0.861 |
| | SpliceBERT | 3435 | 93 | 4958 | 9425 | 0.673 | 0.860 |
| | SSNet_gtex | 3620 | 104 | 4947 | 9240 | 0.679 | 0.863 |
| | SSNet_pangolin | 2913 | 36 | 5015 | 9947 | 0.680 | 0.866 |
| | SSNet_gtex_pangolin | 3070 | 42 | 5009 | 9790 | 0.678 | 0.865 |
| | SSNet_pangolin_gtex | 3668 | 103 | 4948 | 9192 | 0.678 | 0.862 |

Supplementary Table S5: Validation Results for BRCA. TP, FP, TN, FN counts at fixed threshold are provided, along with AUROC and AUPRC.

| Label | Model | #TP | #FP | #TN | #FN | AUROC | AUPRC |
|---|---|---|---|---|---|---|---|
| label_expert | SpliceAI | 317 | 29 | 2470 | 4481 | 0.781 | 0.851 |
| | Pangolin | 517 | 19 | 2485 | 4311 | 0.844 | 0.897 |
| | Spliceformer | 328 | 18 | 2486 | 4581 | 0.796 | 0.855 |
| | SpliceTransformer | 337 | 19 | 2485 | 4567 | 0.599 | 0.773 |
| | SpliceBERT | 582 | 17 | 2487 | 4327 | 0.686 | 0.820 |
| | SSNet_gtex | 231 | 22 | 2482 | 4678 | 0.826 | 0.871 |
| | SSNet_pangolin | 239 | 5 | 2499 | 4670 | 0.904 | 0.929 |
| | SSNet_gtex_pangolin | 247 | 5 | 2499 | 4662 | 0.884 | 0.916 |
| | SSNet_pangolin_gtex | 256 | 26 | 2478 | 4653 | 0.831 | 0.874 |
| label_all | SpliceAI | 869 | 133 | 6397 | 6853 | 0.744 | 0.756 |
| | Pangolin | 1170 | 120 | 6440 | 6607 | 0.813 | 0.820 |
| | Spliceformer | 786 | 114 | 6452 | 7173 | 0.750 | 0.755 |
| | SpliceTransformer | 833 | 77 | 6485 | 6994 | 0.601 | 0.686 |
| | SpliceBERT | 1150 | 74 | 6492 | 6809 | 0.637 | 0.719 |
| | SSNet_gtex | 797 | 67 | 6499 | 7162 | 0.805 | 0.800 |
| | SSNet_pangolin | 769 | 23 | 6543 | 7190 | 0.883 | 0.872 |
| | SSNet_gtex_pangolin | 791 | 28 | 6538 | 7168 | 0.859 | 0.850 |
| | SSNet_pangolin_gtex | 824 | 86 | 6480 | 7135 | 0.809 | 0.802 |
